# Increased EZH2 function in regulatory T cells promotes their capacity to suppress autoimmunity by driving effector differentiation prior to activation

**DOI:** 10.1101/2024.04.05.588284

**Authors:** Janneke G.C. Peeters, Stephanie Silveria, Merve Ozdemir, Srinivas Ramachandran, Michel DuPage

## Abstract

The immunosuppressive function of regulatory T (Treg) cells is essential for maintaining immune homeostasis. Enhancer of zeste homolog 2 (EZH2), a histone H3 lysine 27 (H3K27) methyltransferase, plays a key role in maintaining Treg cell function upon CD28 co-stimulation, and *Ezh2* deletion in Treg cells causes autoimmunity. Here we assessed whether increased EZH2 activity in Treg cells would improve Treg cell function. Using an Ezh2 gain-of-function mutation, *Ezh2^Y641F^*, we found that Treg cells expressing *Ezh2^Y641F^* displayed an increased effector Treg phenotype and were poised for improved homing to organ tissues. Expression of *Ezh2^Y641F^* in Treg cells led to more rapid remission from autoimmunity. H3K27me3 profiling and transcriptomic analysis revealed a redistribution of H3K27me3, which prompted a gene expression profile in naïve *Ezh2^Y641F^* Treg cells that recapitulated aspects of CD28-activated *Ezh2^WT^* Treg cells. Altogether, increased EZH2 activity promotes the differentiation of effector Treg cells that can better suppress autoimmunity.

**Highlights:**

- EZH2 function promotes effector differentiation of Treg cells.
- EZH2 function promotes Treg cell migration to organ tissues.
- EZH2 function in Treg cells improves remission from autoimmunity.
- EZH2 function poises naïve Treg cells to adopt a CD28-activated phenotype.

## Introduction

Regulatory T (Treg) cells are an immunosuppressive subset of CD4^+^ T cells that express the transcription factor Foxp3 and restrain effector T (Teff) cell responses ^1^. Treg cells are critical for maintaining immune homeostasis since loss of Treg cells or loss of Foxp3 expression leads to the development of severe autoimmunity in both humans and mice ^2,3^. The pivotal role of Treg cells observed across a broad range of diseases, such as autoimmune disorders, cancer, infection, and transplantation, makes them a clinically important therapeutic target ^2,4–11^.

For autoimmune diseases and transplantation, Treg cell therapeutic approaches have focused on increasing the number of Treg cells, either by autologous or allogeneic Treg cell transfer ^4,11,12^. However, the discovery of mechanisms to either improve or inhibit Treg cell function *in situ* by the delivery of therapeutics that can alter Treg cell function *in vivo* could be used to treat the two extremes of decreased or increased Treg cell function in autoimmunity and cancer, respectively. Regulators of the epigenome can switch sets of genes on or off at a broad level, altering cellular states that can change Treg cell stability and function ^13^. Foxp3 expression is prominently controlled by DNA (hypo)methylation at key Treg cell-specific demethylated regions (TSDRs) in the Foxp3 promoter and the conserved non-coding DNA sequence 2 (CNS2) enhancer region ^13–16,16–22^. Chromatin-modifying enzymes have also been shown to contribute to Treg cell development and function by directly acting on the TSDRs ^18,23–25^. Enhancer of zeste homolog 2 (Ezh2), a subunit of the polycomb repressive complex 2 (PRC2) responsible for catalyzing (tri)-methylation of lysine 27 of histone H3 to make H3K27me3, has been shown to be essential for Foxp3-driven repression of pro-inflammatory genes in Treg cells ^26^. Deletion of *Ezh2* in Treg cells impaired immune homeostasis, reduced Treg cell stability, and disrupted the Foxp3-driven Treg cell program, demonstrating that *Ezh2* is critical for the maintenance of Treg cell identity ^27,28^. Furthermore, EZH2 function is needed in tumor-infiltrating Treg cells to block anti-tumor T cell responses, and the pharmacological targeting of EZH2 activity with a small molecule inhibitor could selectively reprogram intratumoral Treg cells and promote anti-cancer immunity ^29–31^. Interestingly, Treg cell deletion of the lysine demethylase 6B KDM6B/JMJD3, which opposes EZH2 function by removing methylation marks from H3K27, increased tumor outgrowth, suggesting that broad regulation of H3K27me3 levels in Treg cells can regulate Treg cell function^30^.

To further investigate the impact of increased H3K27me3 levels on Treg cell function, we utilized a hyperactive version of EZH2 identified in follicular lymphomas (FL) and germinal-center B-cell-like (GCB) diffuse large B-cell lymphomas (DLBCL) ^32–37^. This gain-of-function allele of Ezh2 contains a single amino acid substitution from tyrosine to phenylalanine at position 641 within the catalytic SET domain (EZH2^Y641F^), leading to greater H3K27me3 levels in cells expressing EZH2^Y641F 38–44^. Using a Cre/LoxP activated allele of *Ezh2^Y641F^* to induce its expression selectively in Treg cells, we demonstrate that EZH2^Y641F^ expression in Treg cells increased their Foxp3 stability and effector differentiation, as well as improved their competitive advantage over wild-type Treg cells in mice co-populated with *Ezh2^Y641F^* and *Ezh2^WT^* Treg cells. This *Ezh2^Y641F^* Treg cell phenotype was rapidly adopted upon acute activation of the EZH2^Y641F^ allele *in vivo,* and the presence of *Ezh2^Y641F^* Treg cells led to a more rapid reversal of autoimmune disease. Genomic analysis of H3K27me3 modifications and gene expression patterns revealed that *Ezh2^Y641F^* Treg cells adopted features of CD28 activation in their naïve state, thus priming Tregs for effector differentiation and homing to organ tissues. Overall, increased EZH2 activity in Treg cells promoted better control of autoimmunity, suggesting that drugs that boost H3K27me3 levels could be a promising therapeutic approach in the setting of transplantation tolerance and autoimmunity.

## Results

### Treg cells with hyperactive *Ezh2* have increased Foxp3 stability and maintain immune homeostasis

To study the effect of increased H3K27me3 levels in Treg cells *in vivo*, we generated mice expressing an EZH2 SET-domain encoding gain-of-function mutation (Y641F) specifically in Treg cells by generating *Foxp3-GFP-hcre;Ezh2^LSL-Y641F^* (Treg.*Ezh2^Y641F^*) mice (Figure 1A) ^43,45^. We also incorporated a lineage-tracing strategy by including an *R26^LSL-RFP^* allele in mice to distinguish stable Treg cells (CD4^+^GFP^+^RFP^+^) from cells that do not maintain Foxp3 expression (CD4^+^GFP^-^RFP^+^)^27^. All analyses in this study were of Treg cells expressing one *Ezh2^Y641F^* and one wild-type *Ezh2* allele (termed *Ezh2^Y641F^* Treg cells). H3K27me3 levels in *Ezh2^Y641F^* Treg cells from lymph node (LN) or spleen were increased compared to Treg cells expressing two wild-type *Ezh2* alleles (termed *Ezh2^WT^* Treg cells) or CD4 effector T cells (CD4^+^Foxp3^-^) (Figure 1B-1C and S1A-1B) ^42–44^. Treg.*Ezh2^Y641F^* mice exhibited reduced Treg cell frequencies in LN and spleen (Figure 1D and Figure S1C). However, this reduction in Treg cell frequency did not impact the effector T cell compartment since CD4^+^ and CD8^+^ T cell frequencies, activation status, and cytokine production were unchanged (Figure 1E and S1D-1E). This suggests that a smaller proportion of Tregs in Treg.*Ezh2^Y641F^* mice may be sufficient to maintain immune homeostasis. Since deletion of *Ezh2* was shown to reduce Treg cell stability, we tested whether increased *Ezh2* activity would impact Foxp3 stability in Treg cells ^27^. As shown in Figure 1F, the percentage of stable CD4^+^GFP^+^RFP^+^ Treg (as a frequency of CD4^+^RFP^+^) cells was increased in Treg.*Ezh2^Y641F^* mice, while CD4^+^GFP^-^RFP^+^ exTreg cells were reduced (Figure 1F). Thus, a hyperactive allele of *Ezh2* increases H3K27me3 levels and Treg stability in secondary lymphoid organs and maintains immune homeostasis in mice.

**Figure 1.**
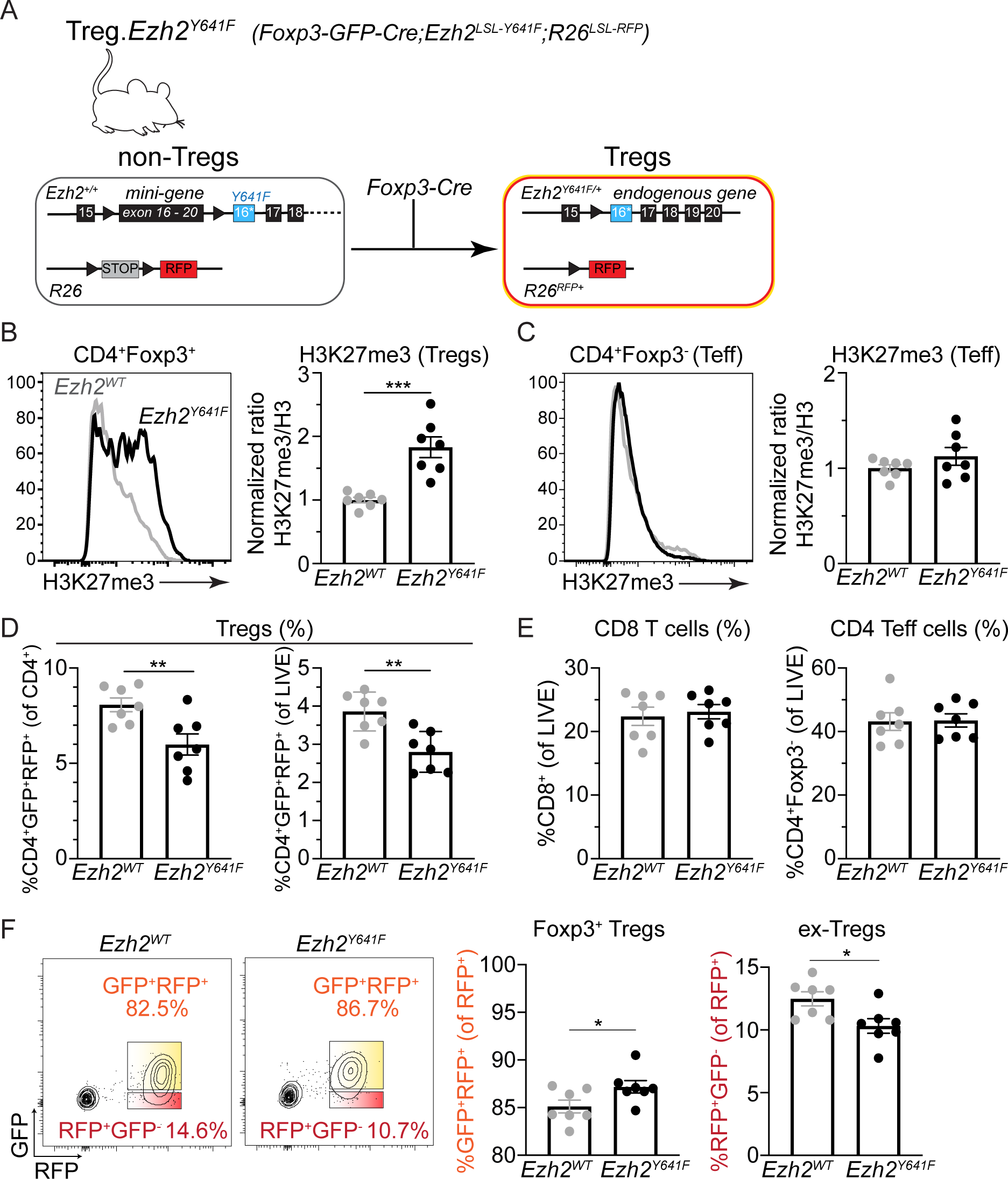
*Ezh2^Y641F^* Treg cells have increased Foxp3 stability and maintain immune homeostasis. (A) Description of alleles used to generate Treg.*Ezh2^Y641F^* mice. (B-C) Representative flow cytometry plots for H3K27me3 staining and normalized H3K27me3/Histone H3 levels in CD4^+^Foxp3^+^ Treg cells (B) or CD4^+^Foxp3^-^ (Teff) cells (C) from LN of Treg.*Ezh2^WT^* (gray lines) and Treg.*Ezh2^Y641F^* (black lines) mice. (A) (D) Frequency of GFP^+^RFP^+^ cells of CD4^+^ and LIVE cells in LN of Treg.*Ezh2^WT^* and Treg.*Ezh2^Y641F^* mice. (B) (E) Frequency of CD8^+^ and CD4^+^ Teff of LIVE cells in LN of Treg.*Ezh2^WT^* and Treg.*Ezh2^Y641F^* mice. (C) (F) Representative flow plots (left) and quantification of the frequencies (right) of GFP^+^RFP^+^ Treg cells and RFP^+^GFP^-^ exTreg cells (as a percentage of CD4^+^RFP^+^ cells) in LN of Treg.*Ezh2^WT^* and Treg.*Ezh2^Y641F^* mice. Data are mean ± SEM and representative of at least two independent experiments (n=12-13 mice total per genotype). Unpaired two-tailed Student’s t-test; *p < 0.05, **p < 0.01, ***p < 0.001, and ****p<0.0001 (only statistically significant differences are noted and non-significant data is not indicated). See also Figure S1.

### *Ezh2^Y641F^* Treg cells exhibit increased activation and effector differentiation

The maintenance of immune homeostasis despite fewer Treg cells in Treg.*Ezh2^Y641F^* mice led us to hypothesize that *Ezh2^Y641F^* Treg cells may exhibit enhanced functionality compared to *Ezh2^WT^* Treg cells. Upon *ex vivo* stimulation, *Ezh2^Y641F^* Treg cells had increased production of the anti-inflammatory cytokine IL-10 (Figure 2A and S2A) ^46–48^. *Ezh2^Y641F^* Treg cells had increased expression of the co-inhibitory markers PD-1 and TIGIT, which are both associated with an effector Treg cell phenotype and are highly expressed on tumor-infiltrating Treg cells (Figure 2B-2C and S2B-C) ^49–56^. The frequency of CD69^+^ Treg cells was also increased, indicating that *Ezh2^Y641F^* Tregs are more activated compared to *Ezh2^WT^* Treg cells. However, the frequency of CD25^+^ Treg cells was decreased in Treg.*Ezh2^Y641F^* mice, which was surprising as it is a key marker of Treg cell identity. Decreased CD25 has been described to occur during *in vivo* proliferation of Treg cells, acting as a negative feedback mechanism that limits Treg cell proliferation ^57,58^. In addition, CD25 has been shown to be less critical for Treg cell maintenance in non-lymphoid organs ^59–61^. CD103 expression on *Ezh2^Y641F^* Treg cells was also increased and has been associated with increased proliferation, an effector memory phenotype, and epithelial tissue homing of Treg cells (Figure 2D and S2D) ^62–66^. CD103 is the integrin alpha E subunit that binds to integrin beta 7 to form the heterodimeric integrin molecule αEβ7, which interacts with E-cadherin, a defining marker of epithelium ^67^. CD103 expression on Tregs has been demonstrated to be important for the migration and retention of Treg cells in epithelial tissues, such as the lung or skin, during homeostasis and inflammation ^62,68,69^. UMAP analysis showed that the expression of these receptors in *Ezh2^Y641F^* Treg cells was sometimes overlapping but also distinct, indicating that increased EZH2 activity leads to a heterogenous Treg phenotype rather than constitutively increasing the expression of all of these receptors in single cells, similar to what is observed with *Ezh2^WT^* Treg cells (Figure 2E and S2E).

**Figure 2.**
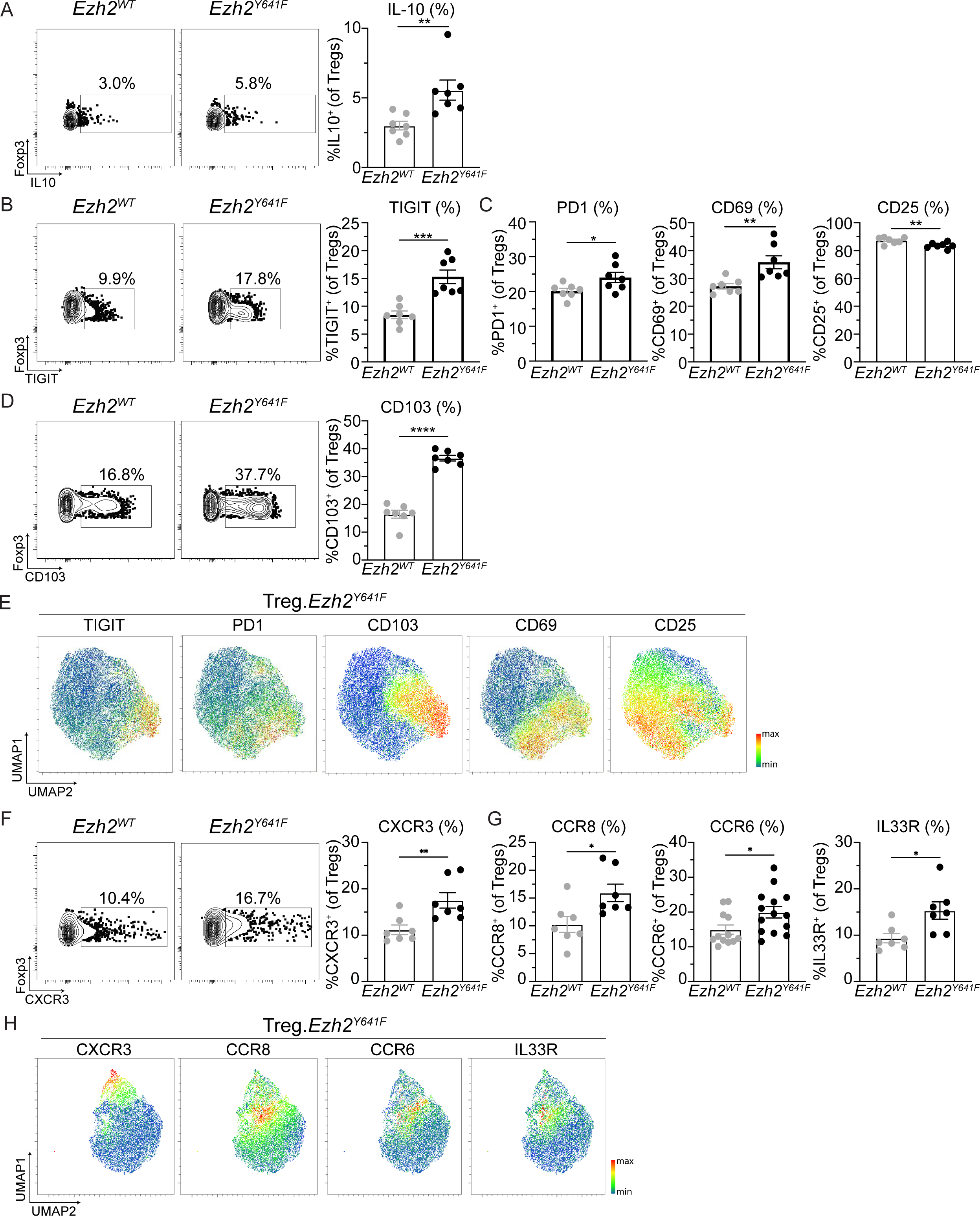
*Ezh2^Y641F^* Treg cells exhibit increased activation and effector differentiation. (A) Representative flow cytometry plots and quantified frequencies of IL-10^+^ Treg cells (CD4^+^Foxp3^+^) from LN of Treg.*Ezh2^WT^* and Treg.*Ezh2^Y641F^* mice. (B) Representative flow cytometry plots and quantified frequencies of TIGIT^+^ Treg cells from LN of Treg.*Ezh2^WT^* and Treg.*Ezh2^Y641F^* mice. (C) Quantified frequencies of Treg cells that are PD1^+^, CD69^+^, or CD25^+^ from LN of Treg.*Ezh2^WT^* and Treg.*Ezh2^Y641F^* mice. (D) Representative flow cytometry plots and quantified frequencies of CD103^+^ Treg cells from LN of Treg.*Ezh2^WT^* and Treg.*Ezh2^Y641F^* mice. (E) UMAP analysis demonstrating the expression of CD103, TIGIT, PD1, CD69, and CD25 in Treg cells from LN of Treg.*Ezh2^Y641F^* mice. (F) Representative flow cytometry plots and quantified frequencies of CXCR3^+^ Treg cells from LN of Treg.*Ezh2^WT^* and Treg.*Ezh2^Y641F^* mice. (G) Quantified frequencies of Treg cells that are CCR8^+^, CCR6^+^, or IL33R^+^ from LN of Treg.*Ezh2^WT^* and Treg.*Ezh2^Y641F^* mice. (H) UMAP analysis demonstrating the expression of CXCR3, CCR8, CCR6, and IL33R in Treg cells from LN of Treg.*Ezh2^Y641F^* mice. Data are mean ± SEM and representative of at least two independent experiments (n=12-13 mice per genotype). Unpaired two-tailed Student’s t-test; *p < 0.05, **p < 0.01, ***p < 0.001, and ****p<0.0001 (only statistically significant differences are noted and non-significant data is not indicated). See also Figure S2.

Treg cell migration and retention in non-lymphoid tissues is also regulated by the expression of chemokine receptors. Analysis of CXCR3, CCR8, and CCR6, each defining chemokine receptors in varying Th-like Treg cell responses, revealed that *Ezh2^Y641F^* compared to *Ezh2^WT^* Treg cells had increased expression of each, indicating that *Ezh2^Y641F^* Treg cells may be prone to differentiating into multiple Th-like Treg phenotypes and migrating to non-lymphoid organ tissues (Figure 2F-G and S2F-G) ^70–74^. In addition, expression of the ST2/IL33 receptor was increased on *Ezh2^Y641F^* compared to *Ezh2^WT^* Treg cells, and IL33R expression is associated with wound healing and repair after tissue damage ^75–78^. UMAP analysis clearly showed distinct patterns of expression of each of these receptors in *Ezh2^Y641F^* Treg cells, suggesting again that the *Ezh2^Y641F^* Treg cells generate a heterogenous population of effector Treg cells rather than a generalized, multipotent Treg phenotype, which is similar to *Ezh2^WT^* Treg cells (Figure 2H and S2H). These characteristics of enhanced function and effector differentiation of lymphoid Treg cells from Treg.*Ezh2^Y641F^* mice may underlie the maintenance of immune homeostasis in mice despite fewer Treg cells. Furthermore, *Ezh2^Y641F^* expression appears to endow Treg cells with a differentiated state that could promote enhanced migration to organ tissues and sites of inflammation.

### *Ezh2^Y641F^* Treg cells have a competitive advantage over WT Treg cells

*Ezh2^Y641F^* Treg cells exhibited increased activation and effector differentiation compared to *Ezh2^WT^* Treg cells. Therefore, we tested whether *Ezh2^Y641F^* Treg cells would be at a competitive advantage in mice co-populated with *Ezh2^WT^* Treg cells. Using *Foxp3^GFP-DTR^/Foxp3^YFP-Cre^;Ezh2^LSL-Y641F/+^;R26^LSL-RFP^* female mice (hereafter referred to as Treg.*Ezh2^Y641F^/Ezh2^WT^* mice), we analyzed the contribution of *Ezh2^Y641F^* versus *Ezh2^WT^* Treg cells to the total pool of Treg cells ^6,27,30,79^. Since *Foxp3* is located on the X chromosome, which undergoes random X inactivation, female mice with one *Foxp3^YFP-Cre^* allele will express YFP-Cre protein in approximately half of their Treg cells, while the rest of the Treg cells will express DTR-GFP from the other *Foxp3^GFP-DTR^* allele. In combination with the *Ezh2^LSL-Y641F^* and *R26 ^LSL-RFP^* alleles, half of the Treg cells will express *Ezh2^Y641F^* and be identifiable by RFP expression, whereas RFP^-^ Treg cells will express *Ezh2^WT^* (Figure 3A). As a control, *Foxp3^GFP-DTR^/Foxp3^YFP-Cre^;Ezh2^+/+^;R26^LSL-RFP^* mice (Treg.*Ezh2^WT^/Ezh2^WT^* mice) were analyzed, as we have observed a negative impact of YFP-Cre compared to DTR-GFP expression in Treg cells in this competitive setting (see Figure 3B, which shows that Treg frequencies are not 50:50). Comparison of the distribution between *Ezh2^Y641F^* Treg cells (YFP^+^RFP^+^) and WT Treg cells (GFP^+^RFP^-^) revealed that the ratio YFP^+^RFP^+^/GFP^+^RFP^-^ Treg cells was significantly increased in Treg.*Ezh2^Y641^/Ezh2^WT^* mice compared to mice having only *Ezh2^WT^* Treg cells, indicating that expression of *Ezh2^Y641F^* gives Treg cells a competitive advantage in the lymph node and spleen (Figure 3B and S3A). A direct comparison of H3K27me3 levels between *Ezh2^Y641F^* and *Ezh2^WT^* Treg cells confirmed that H3K27me3 was increased in *Ezh2^Y641F^* Treg cells (Figure 3C and Figure S3B). However, CD4^+^ and CD8^+^ T effector cell frequencies, activation status, or cytokine production, as well as the frequency of total Treg cells, were not changed in Treg.*Ezh2^Y641^/Ezh2^WT^* mice compared to Treg.*Ezh2^WT^/Ezh2^WT^* mice (Figure 3D and Figure S3C-3D).

**Figure 3.**
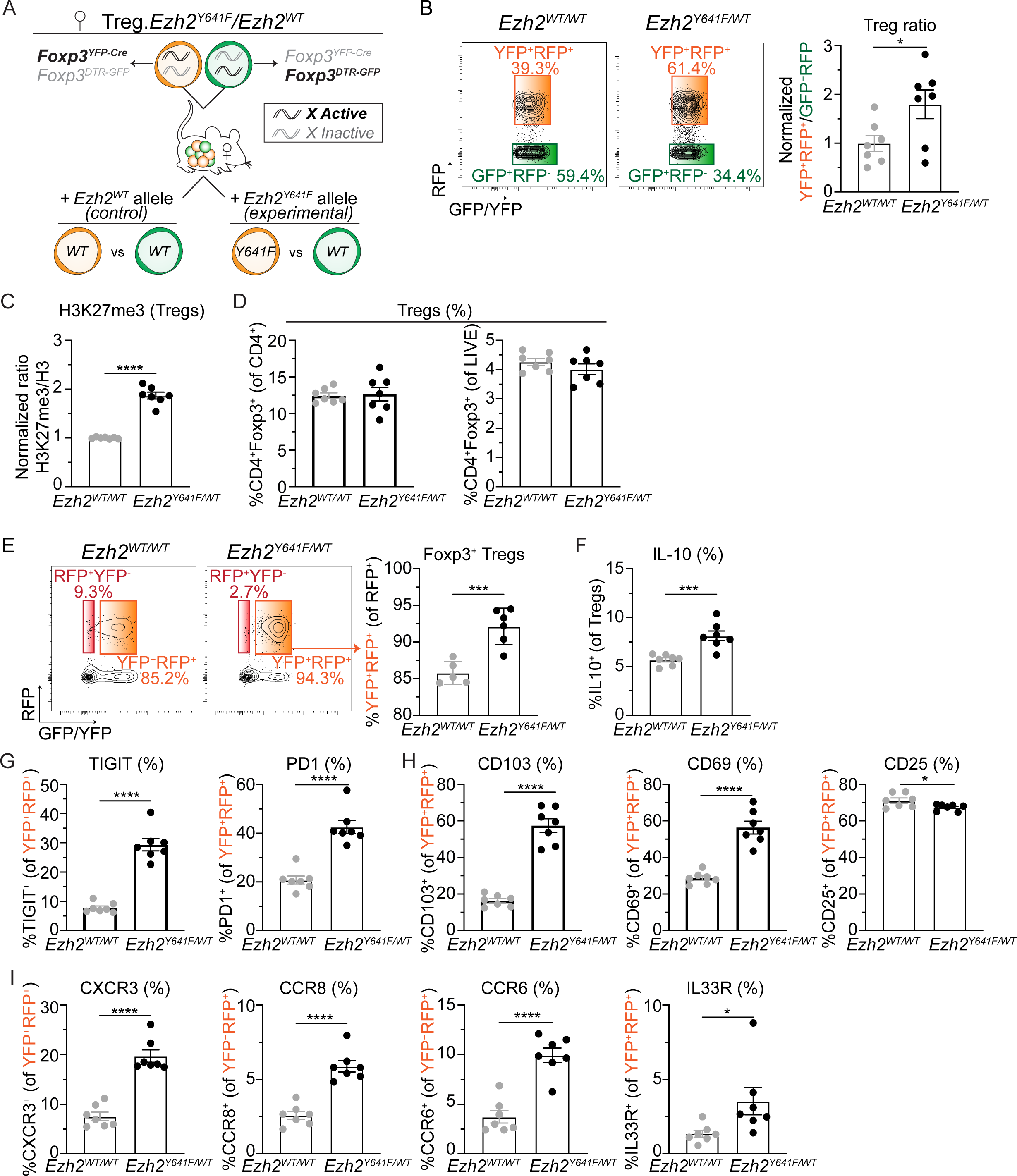
*Ezh2^Y641F^* Treg cells have a competitive advantage over WT Treg cells. (A) Schematic model of female mosaic mice used to study Treg cells in a competitive setting. (B) Representative flow cytometry plots and normalized ratios of YFP^+^RFP^+^/GFP^+^RFP^-^ Treg cells quantified from LN of Treg.*Ezh2^WT^/Ezh2^WT^* compared to Treg.*Ezh2^Y641^/Ezh2^WT^* mice. (C) Normalized H3K27me3/Histone H3 levels of CD4^+^Foxp3^+^ Treg cells in LN of Treg.*Ezh2^WT^/Ezh2^WT^* compared to Treg.*Ezh2^Y641^/Ezh2^WT^* mice. (D) Quantified frequencies of CD4^+^Foxp3^+^ of CD4^+^ and LIVE cells in LN of Treg.*Ezh2^WT^/Ezh2^WT^* and Treg.*Ezh2^Y641^/Ezh2^WT^* mice. (E) Representative flow cytometry plots and quantified frequencies of YFP^+^RFP^+^ (as a percentage of CD4^+^RFP^+^ cells) in LN of Treg.*Ezh2^WT^/Ezh2^WT^* and Treg.*Ezh2^Y641^/Ezh2^WT^* mice. (F) Quantified frequencies of IL-10^+^ Treg cells in LN of Treg.*Ezh2^WT^/Ezh2^WT^* and Treg.*Ezh2^Y641^/Ezh2^WT^* mice. (G) Quantified frequencies of TIGIT^+^ and PD1^+^ of YFP^+^RFP^+^ Treg cells in LN of Treg.*Ezh2^WT^/Ezh2^WT^* and Treg.*Ezh2^Y641^/Ezh2^WT^* mice. (H) Quantified frequencies of PD1^+^, CD103^+^, CD69^+^, CD25^+^ of YFP^+^RFP^+^ Treg cells in LN of Treg.*Ezh2^WT^/Ezh2^WT^* and Treg.*Ezh2^Y641^/Ezh2^WT^* mice. (I) Quantified frequencies of CXCR3^+^, CCR8^+^, CCR6^+^, and IL33R^+^ of YFP^+^RFP^+^ Treg cells in LN of Treg.*Ezh2^WT^/Ezh2^WT^* and Treg.*Ezh2^Y641^/Ezh2^WT^* mice. Data are mean ± SEM and representative of at least two independent experiments (n=8-11 mice total per genotype). Unpaired two-tailed Student’s t-test; *p < 0.05, **p < 0.01, ***p < 0.001, and ****p<0.0001 (only statistically significant differences are noted and non-significant data is not indicated). See also Figure S3.

As observed in *Foxp3-GFP-hcre;Ezh2^LSL-Y641F/+^* mice, Foxp3 stability and IL-10 production were enhanced in *Ezh2^Y641F^* Treg cells in mice co-populated with *Ezh2^WT^* Treg cells, indicating the intrinsic activity of *Ezh2^Y641F^* on the Treg cell phenotype (Figure 3E-3F and S3E). Furthermore, analysis of co-inhibitory, activation, migration, and differentiation markers showed the same changes in expression observed in *Ezh2^Y641F^* Treg cells from *Foxp3-GFP-hcre;Ezh2^LSL-Y641F/+^* mice (Figure 3G-I and Figure S3F-3H). In summary, increased EZH2 activity in Treg cells increases Foxp3 stability as well as their activation and effector differentiation state intrinsically, leading *Ezh2^Y641F^* Treg cells to exhibit a competitive advantage over WT Treg cells when both cell types populate mice.

### Acute induction of *Ezh2^Y641F^* in Treg cells leads to their rapid adoption of an activated and effector differentiated phenotype

To distinguish whether hyperactivation of EZH2 impacts Treg cell phenotypes due to EZH2 activity during Treg cell development or if EZH2 hyperactivation in mature Treg cells post-development also induces similar changes, we used a tamoxifen-inducible CreER allele to activate *Ezh2^Y641F^* in differentiated Treg cells *in vivo* ^80^. Using *Foxp3^eGFP-Cre-ERT2^;Ezh2^Y641F/+^* mice (hereafter referred to as Treg.*iEzh2^Y641F^*), we induced the expression of *Ezh2^Y641F^* by tamoxifen administration and assessed Treg cells two weeks later (Figure 4A) ^30^. Analysis of H3K27me3 levels in Treg cells from these mice compared to Treg cells from Treg.*iEzh2^WT^* revealed increased H3K27me3 levels in Treg.*iEzh2^Y641F^* mice but no change in the total frequencies of Treg cells (Figure 4B-4C and S4A-S4B). Based on PD1, TIGIT, CD69, CD103, and CD25 expression, *iEzh2^Y641F^* Treg cells already displayed increased effector differentiation at this time point (Figure 4D-4E). Analysis of the different chemokine receptors further confirmed that within two weeks of the expression of hyperactive *Ezh2,* Treg cells acquired a heterogeneous effector differentiated phenotype (Figure 4F). However, the intrinsic suppressive activity of Treg cells *in vitro* was not affected (Figure S4C). As previously observed in the presence of constitutive *Ezh2^Y641F^* Treg cells, CD4^+^ and CD8^+^ T effector cell frequencies, activation status, and cytokine production were not changed in tamoxifen-treated Treg.*iEzh2^Y641F^* mice compared to Treg.*iEzh2^WT^* mice (Figure S4D). The lack of a reduction in Treg cell frequency with inducible EZH2 hyperactivation in Treg.*iEzh2^Y641F^* mice (Figure 4C) was different than what we observed in *Foxp3-GFP-hcre;Ezh2^Y641F/+^* mice (Figure 1D), wherein Treg cells had constitutively hyperactive EZH2 and *Ezh2^Y641F^* Treg cell frequencies were reduced. We hypothesize that the reduction in Treg cells in mice constitutively expressing *Ezh2^Y641F^* was an indirect effect of *Ezh2^Y641F^* Treg cells being more functional, thus allowing for fewer Treg cells to maintain immune homeostasis over a prolonged period of time. In the setting of acute *Ezh2^Y641F^* activation, the rebalancing of Treg frequencies required to maintain immune homeostasis has not had enough time to recalibrate. Overall, inducible expression of *Ezh2^Y641F^* led to the rapid adoption of an activated and effector differentiated Treg cell phenotype similar to constitutive *Ezh2^Y641F^* expression in Treg cells. This supports the hypothesis that hyperactive EZH2 rapidly impacts the phenotype of mature Treg cells *in vivo* in a cell-intrinsic manner.

**Figure 4.**
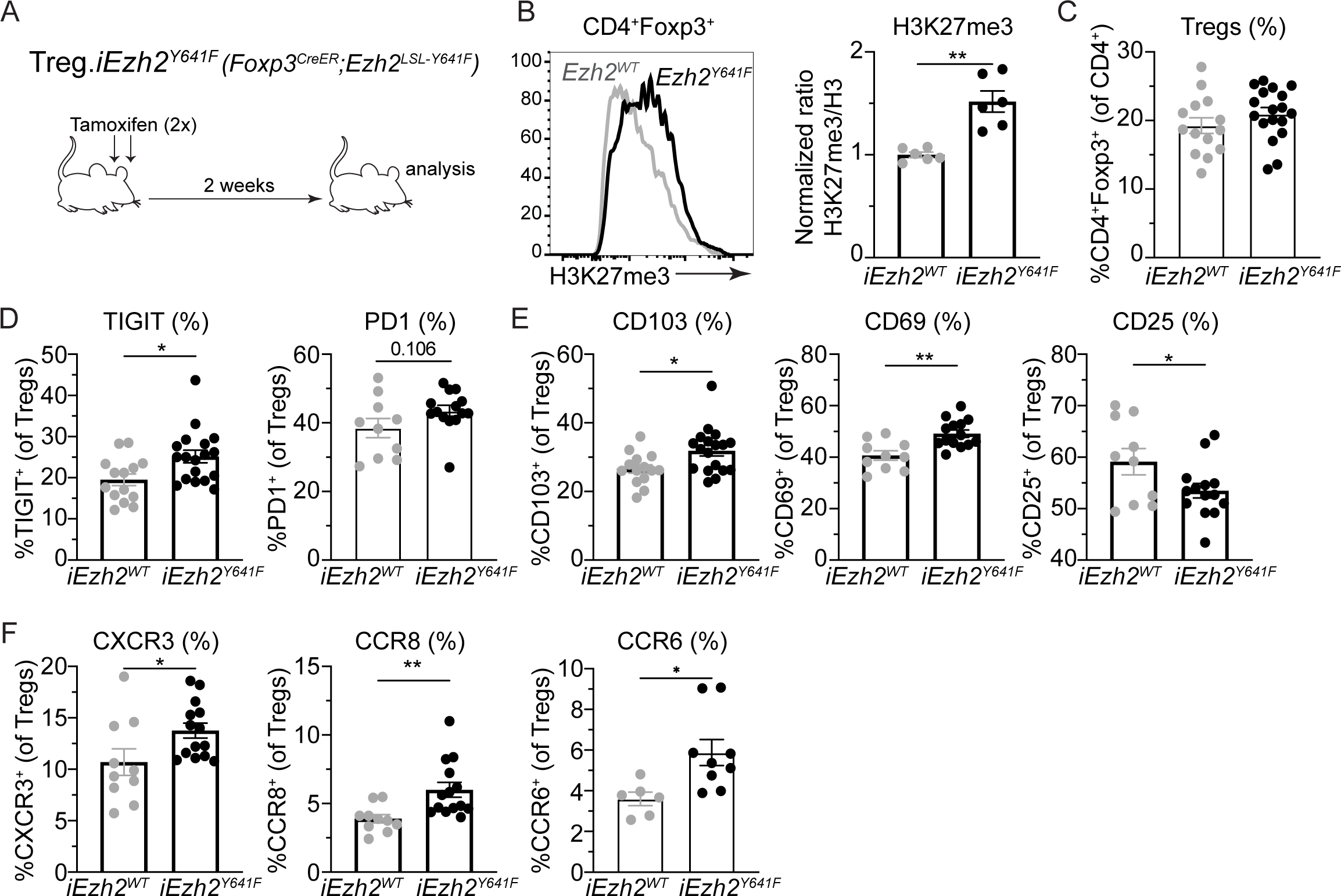
Acute induction of *Ezh2^Y641F^* in Treg cells increases their activation and effector differentiation state. (A) Schematic depiction of the tamoxifen-inducible activation of the *Ezh2^Y641F^* allele in Treg cells. (B) Normalized H3K27me3/Histone H3 levels of CD4^+^Foxp3^+^ Treg cells in LN of Treg.*iEzh2^WT^* compared to Treg.*iEzh2^Y641F^* mice treated as depicted in A. (C) Quantified frequencies of CD4^+^Foxp3^+^ Treg cells in LN of Treg.*iEzh2^WT^* and Treg.*iEzh2^Y641F^* mice treated as depicted in A. (D) Quantified frequencies of TIGIT^+^ and PD1^+^ Treg cells in LN of Treg.*iEzh2^WT^* and Treg.*iEzh2^Y641F^* mice treated as depicted in A. (E) Quantified frequencies of CD103^+^, CD69^+^, and CD25^+^ Treg cells in LN of Treg.*iEzh2^WT^* and Treg.*iEzh2^Y641F^* mice treated as depicted in A. (F) Quantified frequencies of CXCR3^+^, CCR8^+^, and CCR6^+^ Treg cells in LN of Treg.*iEzh2^WT^* and Treg.*iEzh2^Y641F^* mice treated as depicted in A. Data are mean ± SEM (n=6-18 mice per group pooled from two independent experiments). Unpaired two-tailed Student’s t-test; *p < 0.05, **p < 0.01, ***p < 0.001, and ****p<0.0001 (only statistically significant differences are noted and non-significant data is not indicated). See also Figure S4.

### *Ezh2^Y641F^* Treg cells robustly reverse autoimmunity and outcompete WT Tregs for infiltration in organ tissues

Since inducible expression of *Ezh2^Y641F^* rapidly increased multiple phenotypes in Treg cells associated with increased activation, differentiation, and migration without reducing Treg cell frequencies (Figure 4C-4F), we used this genetic model to explore whether increased *Ezh2* activity in Treg cells could better control autoimmunity. To do so, we made use of the experimental autoimmune encephalomyelitis (EAE) mouse model of multiple sclerosis, where Treg.*iEzh2^Y641F^* and Treg.*iEzh2^WT^* mice were immunized with myelin oligodendrocyte glycoprotein (MOG) in complete Freund’s adjuvant emulsion 2 weeks after tamoxifen treatment and disease score was followed over time. Mice containing *iEzh2^WT^* Tregs and *iEzh2^Y641F^* became equally sick, but mice expressing *iEzh2^Y641F^* recovered significantly better from the disease (Figure 5A). In addition, there were increased MOG/H-2IA^b+^-specific Treg cells in the CNS of Treg.*iEzh2^Y641F^* mice compared to Treg.*iEzh2^WT^* mice (Figure 5B). Our findings are in agreement with previous results showing that Treg cells mediate recovery from EAE by controlling effector T cell proliferation within the CNS ^81–84^. Interestingly, *Ezh2^Y641F^* Treg cells in the draining lymph nodes of mice undergoing EAE displayed an enhanced effector phenotype compared to *Ezh2^WT^* Treg cells, as was observed under homeostatic conditions; however, the phenotype of *Ezh2^Y641F^* Treg cells in the CNS was similar to *iEzh2^WT^* Treg cells (Figure S5A). This suggests that the phenotypic characteristics of *Ezh2^Y641F^* expression may play an important role in priming Treg cell responses in the lymphoid organs for more rapid differentiation and recruitment to organ tissues rather than enhancing their activity as effector Treg cells in organ tissues.

**Figure 5.**
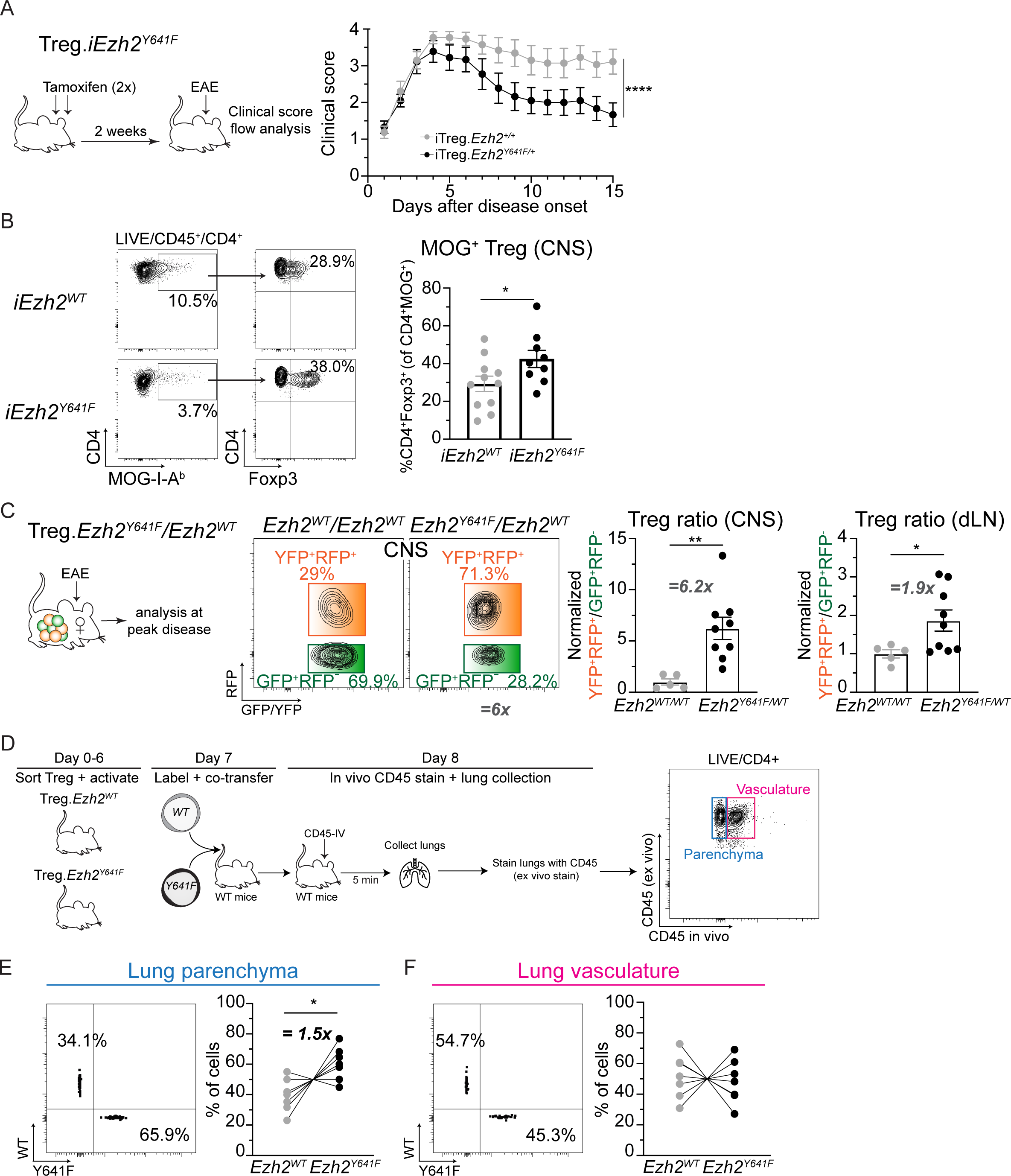
*Ezh2^Y641F^* Treg cells reduce autoimmunity and outcompete WT Tregs for infiltration into organ tissues. (A) Experimental strategy and EAE disease score plotted from time of symptom onset in *Ezh2^Y64F1^* inducible Treg.*iEzh2^WT^* compared to Treg.*iEzh2^Y641F^* mice (n=9-13, pooled from two independent experiments). (B) Representative flow cytometry plots and quantified frequencies of CD4^+^Foxp3^+^ Treg cells of MOG-specific (MOG-I-A^b+^) CD4^+^ T cells in CNS of Treg.*iEzh2^WT^* and Treg.*iEzh2^Y641F^* mice with fulminant EAE disease. (C) Representative flow cytometry plots and normalized ratio YFP^+^RFP^+^/GFP^+^RFP^-^ of Treg cells in CNS and draining LN (dLN) of female mosaic Treg.*Ezh2^WT^/Ezh2^WT^* and Treg.*Ezh2^Y641F^/Ezh2^WT^* mice with fulminant EAE disease. (D) Experimental strategy describing the activation, fluorescent dye-labeling, and co-transfer of *Ezh2^WT^* and *Ezh2^Y641F^* Treg cells into WT mice for localization analysis into the lung. IV-injected anti-CD45 antibody was used to distinguish cells in the vasculature versus the parenchyma of the lung. (E) Representative flow cytometry plot and quantified frequencies of *Ezh2^WT^* and *Ezh2^Y641F^* Treg cells recovered from the lung parenchyma after adoptive transfer. (F) Representative flow cytometry plot and quantified frequencies of *Ezh2^WT^* and *Ezh2^Y641F^* Treg cells recovered from the lung vasculature after adoptive transfer. Data are mean ± SEM (n=7-13 mice per group pooled from two or three independent experiments). Two-way repeated measured ANOVA (A) and unpaired two-tailed Student’s t-test (B, C, E, F) were used; *p < 0.05, **p < 0.01, ***p < 0.001, and ****p<0.0001 (only statistically significant differences are noted and non-significant data is not indicated). See also Figure S5.

To test whether *Ezh2^Y641F^* Treg cells are better poised to migrate to the CNS during autoimmune inflammation, we again used female Treg.*Ezh2^Y641F^/Ezh2^WT^* mice co-populated with *Ezh2^Y641F^* and *Ezh2^WT^* Treg cells. Analysis of the ratio of YFP^+^RFP^+^/GFP^+^RFP^-^ in the CNS of Treg.*Ezh2^Y641F^/Ezh2^WT^* and Treg.*Ezh2^WT^/Ezh2^WT^* mice showed dramatic favoritism for YFP^+^RFP^+^ (*Ezh2^Y641^* Treg) cells over GFP^+^RFP^-^ (*Ezh2^WT^* Treg) cells in mice undergoing EAE (Figure 5C and S5B). This supports the hypothesis that *Ezh2^Y641F^* Treg cells can outcompete WT Tregs for migration into the inflamed CNS but does not rule out better maintenance or survival in the CNS tissue. To directly test whether *Ezh2^Y641F^* Treg cells can migrate better to organ tissues *in vivo*, we adoptively transferred equal numbers of *Ezh2^WT^* and *Ezh2^Y641F^* Treg cells that were FACS purified from Treg.*Ezh2^WT^* or Treg.*Ezh2^Y641F^* mice which were labeled with distinct fluorescent tracking dyes, into healthy wild-type mice (Figure 5D and S5C). After 24 hours, we injected anti-CD45 antibody intravenously five minutes prior to euthanasia to distinguish blood (vasculature) versus tissue-infiltrating (parenchyma) cells (Figure 5D). We then analyzed the lung and spleen for the presence of transferred Treg cells. This analysis revealed that the *Ezh2^Y641F^* Treg cells recovered from the lung parenchyma specifically were increased compared to *Ezh2^WT^* Treg cells (Figure 5E). Notably, the fraction of *Ezh2^Y641F^* Treg cells recovered from the lung blood vasculature or the spleen was not increased compared to *Ezh2^WT^* Treg cells (Figure 5F and S5D-S5E), demonstrating that *Ezh2^Y641F^* Treg cells have an increased capacity to extravasate into the lung parenchyma, the first tissue encountered upon intravenous transfer. Together, this data demonstrates that the activation and effector differentiation characteristics of *Ezh2^Y64F1^* Treg cells endow these Treg cells with an improved capacity to enter organ tissues, potentially improving their ability to control autoimmunity.

### Expression of *Ezh2^Y641F^* leads to a global redistribution of H3K27me3 in Treg cells

To reveal the underlying mechanisms driving the altered phenotype of *Ezh2^Y641F^* Treg cells, we performed H3K27me3 CUT&RUN on naïve and *in vitro*-activated *Ezh2^Y641F^* Treg, *Ezh2^WT^* Treg, and Foxp3^-^CD4^+^ T (Teff) cells. In agreement with our flow cytometric analysis showing that *Ezh2^Y641F^* Treg cells have increased H3K27me3 levels globally (Figure 1B), global levels of H3K27me3 by CUT&RUN signal across the genome also showed that both naïve and activated *Ezh2^Y641F^* Treg cells exhibited increased H3K27me3 compared to *Ezh2^WT^* Treg cells and effector T cells (Figure 6A). Principal component analyses (PCA) for the enrichment of H3K27me3 at gene bodies or gene promoters showed that PC1, which accounted for 85-89% of the variation, captured activation of Teff cells, whereas PC2, which accounted for 5-8% of the variation captured the differences due to the genotype of *Ezh2^Y641F^* Treg versus *Ezh2^WT^* Treg cells (Figure 6B). Thus, the PCA analysis demonstrates that the H3K27me3 enrichment associated with genes can distinguish *Ezh2^Y641F^* and *Ezh2^WT^* Treg cells. The cumulative distribution of H3K27me3 modifications across all gene bodies, from their transcriptional start site (TSS) to the transcription end site (TES), revealed a marked increase in the H3K27me3 signal across the gene body of *Ezh2^Y641F^* Treg cells, particularly in naïve *Ezh2^Y641F^* Treg cells compared to naïve *Ezh2^WT^* Treg cells (Figure 6C). However, the characteristic enrichment of H3K27me3 at the TSS in *Ezh2^WT^* Treg cells was absent in naïve, or decreased in activated, *Ezh2^Y641F^* Treg cells (Figure 6C). The distribution of H2K27me3 peaks across genic features showed that the fraction of peaks overlapping with promoter/TSS regions was decreased in naïve *Ezh2^Y641F^* Treg cells, but the fraction of peaks within intergenic regions was increased (Figure 6D). This suggests that the expression of *Ezh2^Y641F^* in naïve Treg cells leads to a global re-distribution of H3K27me3 modifications from genic to intergenic regions (Figure 6D). Notably, a global redistribution of H3K27me3 modifications in cells expressing *Ezh2^Y641F^* has been described in transformed B cell lymphomas as well as embryonic stem cells ^43,85^. Therefore, although the EZH2^Y641F^ mutation increases EZH2 activity and increases H3K27me3 modifications globally in *Ezh2^Y641F^* Treg cells, it does so while re-distributing the abundance of H3K27me3 away from promoter/TSS regions and toward intergenic regions, rendering the promoters of genes in *Ezh2^Y641F^* Treg cells with significantly reduced H3K27me3 enrichment compared to *Ezh2^WT^* Treg cells (Figure 6E and 6F). Interestingly, the genes associated with decreased H3K27me3 in their promoter regions were most enriched in embryonic and neuronal development gene ontology phenotypes, indicative of the role of EZH2 activity and H3K27me3 modifications in repressing alternate cell states (Figure S6A and S6B) ^86,87^. However, this did not inform the mechanisms of increased effector differentiation of *Ezh2^Y641F^* Treg cells.

**Figure 6.**
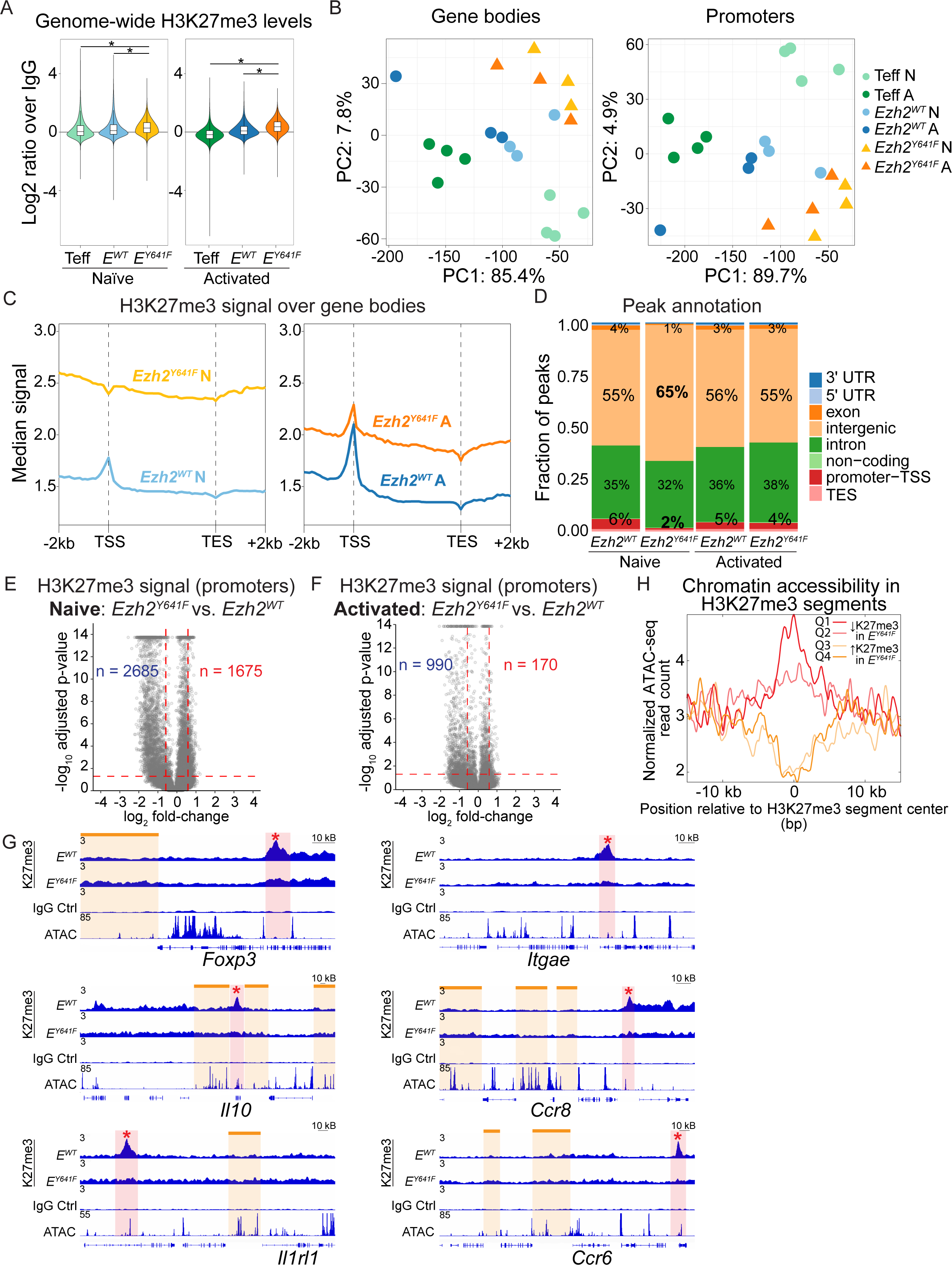
Expression of *Ezh2^Y641F^* leads to a global redistribution of H3K27me3 in Treg cells. (A) Log2 ratio of H3K27me3 signal over IgG across the entire genome for each sample. (B) Principal component analysis based on the top 10,274 variable H3K27me3 regions in gene bodies (left) or the top 9,192 variable H3K27me3 regions in gene promoters (right). (C) Absolute normalized H3K27me3 signal over all gene bodies -/+ 2 kb genome-wide. (D) Fraction of peaks overlapping with genic features. (E-F) Pairwise comparison of H3K27me3 signal in gene promoters in naïve (E) or activated (F) *Ezh2^Y641F^* Treg cells compared to naïve or activated *Ezh2^WT^* Treg cells, respectively. (G) H3K27me3 modification tracks in naïve *Ezh2^WT^* versus *Ezh2^Y641F^* Treg cells and IgG control, as well as representative chromatin accessibility tracks from ATAC-seq data (obtained from GSE233902), for genomic regions surrounding the *Foxp3, Il10, Il1rl1, Itgae, Ccr8,* and *Ccr6* loci. H3K27me3 peaks present only in *Ezh2^WT^* Treg cells (red asterisk) and broadly increased H3K27me3 levels within intergenic regions in *Ezh2^Y641F^* Treg cells (orange lines) are labeled. (H) Metaplot of normalized ATAC-seq read counts centered around H3K27me3 segments with a ≥ 2-fold change in H3K27me3 between naïve *Ezh2^Y641F^* and *Ezh2^WT^* Treg cells and divided into four quartiles (Q1-Q4 defined in Figure S6D). Data are obtained from 3 biological replicates per group. Ordinary two-way ANOVA (A) and Wald test (E and F) were used. *p < 0.05 (only statistically significant differences are noted and non-significant data is not indicated). See also Figure S6.

Therefore, we examined H3K27me3 modifications at the genes we had already identified as differentially induced in *Ezh2^Y641F^* Treg cells from our previous flow cytometric analysis. Interestingly, an examination of the *Foxp3*, *Itgae*, *Tigit*, *Il10*, *Ccr8*, *Ccr6*, and *Il1rl1* genomic loci revealed: (1) the loss of a prominent H3K27me3 peak within 20-100Kb of each gene and (2) an increased deposition of H3K27me3 in adjoining intergenic regions in naïve *Ezh2^Y641F^* compared to *Ezh2^WT^* Treg cells (Figure 6G and S6C). Although a clear role for the large nearby H3K27me3 peaks could not be immediately established, we did find these peaks to lie in regions of accessible chromatin, which is suggestive of nucleation sites for PRC2 (Figure 6G) ^88,89^. To determine whether these lost H3K27me3 peaks in *Ezh2^Y641F^* Treg cells were globally associated with accessible chromatin, we identified H3K27me3-associated DNA segments (as defined in Figure 7A and S6D) that exhibited a ≥ 2-fold change in H3K27me3 between naïve *Ezh2^Y641F^* and *Ezh2^WT^* Treg cells. These H3K27me3 segments (11,696) were then ordered by their fold change in H3K27me3 levels into four quartiles (2,924 each) comparing *Ezh2^Y641F^* and *Ezh2^WT^* Treg cells (Q1 segments had the greatest loss in H3K27me3 in naïve *Ezh2^Y641F^* Treg cells, whereas Q4 segments had the greatest gain in H3K27me3). Overlap with ATAC-sequencing data (GSE233902) from unstimulated Treg cells indicated that H3K27me3 segments in Q1 and Q2 were enriched in ATAC-seq reads at the center of H3K27me3 segments, whereas Q3 and Q4 had a significant depletion of ATAC-seq reads (Figure 6H). Furthermore, the number of overlapping ATAC-seq peaks and H3K27me3 segments was greatly increased in Q1 and Q2 compared to Q3 and Q4, again suggesting that the large H3K27me3 peaks that were lost in naïve *Ezh2^Y641F^* Treg cells occurred in genomic regions associated with highly accessible chromatin across the genome, which is consistent with these being nucleation sites for PRC2 (Figure S6E) ^88,89^. We hypothesize that the reduction in prominent H3K27me3 peaks, along with a concomitant increase in intergenic H3K27me3 levels, leads to reduced H3K27me3-mediated gene repression in naïve *Ezh2^Y641F^* Treg cells. Therefore, enhanced activity of EZH2 may paradoxically lead to increased expression of many genes in *Ezh2^Y641F^* Treg cells to drive their effector differentiation phenotype.

**Figure 7.**
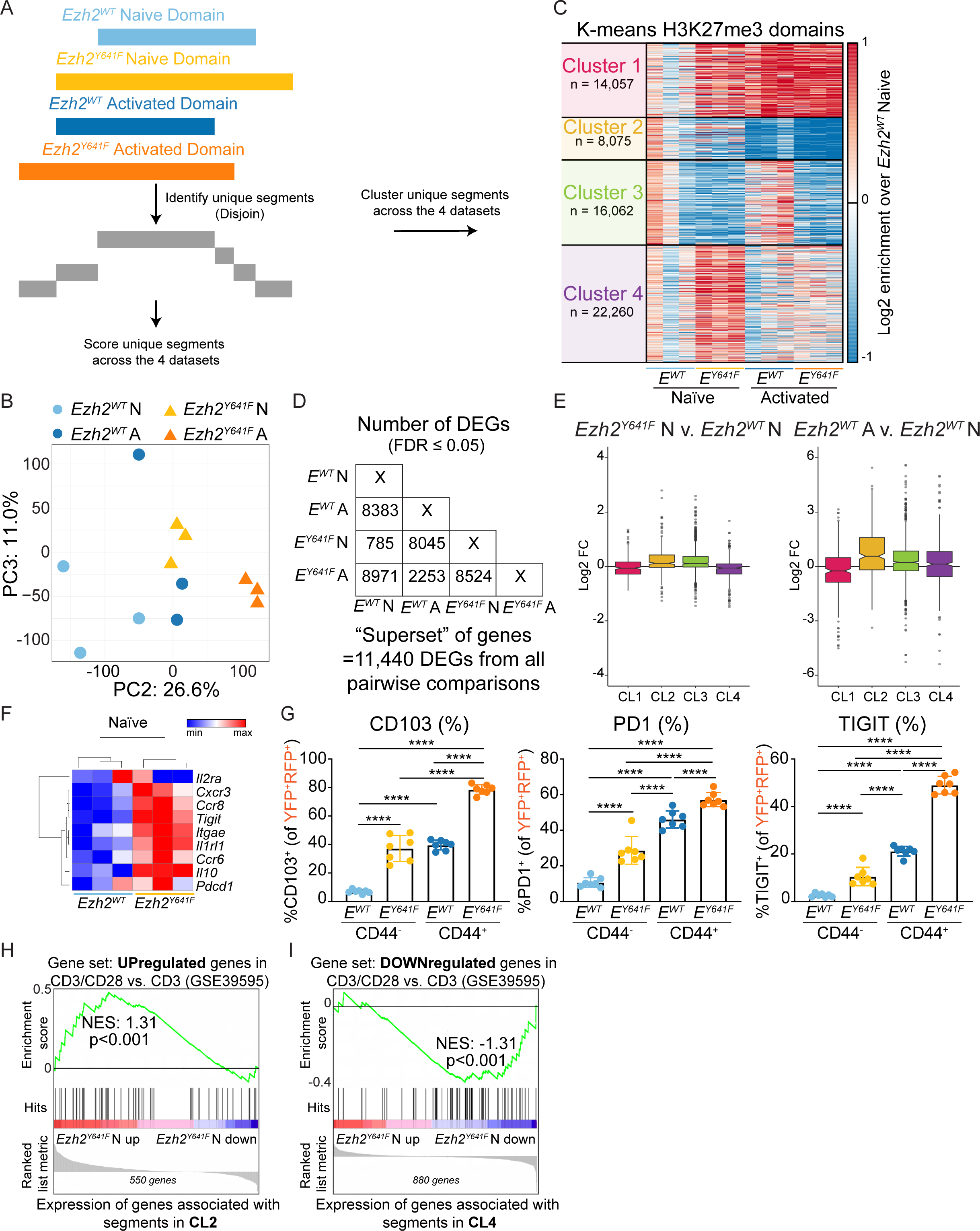
H3K27me3 modifications in naïve *Ezh2^Y641F^* Treg cells drives a gene expression pattern that mimics CD28-activated *Ezh2^WT^* Treg cells. (A) Strategy to define unique, non-overlapping segments by comparison of H3K27me3 domains between naïve and activated, *Ezh2^WT^* and *Ezh2^Y641F^* Treg cells. (B) Principal component analysis of H3K27me3 enrichment across all unique segments. (C) Heatmap of log2 enrichment of H3K27me3 for each dataset over naïve *Ezh2^WT^* dataset at unique segments ordered based on k-means clustering (k=4). (D) Gene expression was compared to generate a list of differentially expressed genes (DEGs) across all six comparisons. A superset of genes (11,440 total), representing an accumulation of DEGs across any pairwise comparison between two groups, was used for correlative gene expression analysis with H3K27me3 levels in E. (E) Boxplots demonstrating the fold change in gene expression between naïve *Ezh2^Y641F^* and *Ezh2^WT^* Treg cells (left) and activated *Ezh2^WT^* and naïve *Ezh2^WT^* Treg cells (right) of genes in the superset (defined in 7D) whose TSS overlapped with H3K27me3 segments in each cluster (CL1-CL4). Statistics for left boxplot is as follows: CL1*; CL2****; CL3****; CL4****, statistics for right boxplot is as follows: CL1***; CL2****; CL3****; CL4**. (F) Heatmap depicting normalized gene expression of selected genes in naïve *Ezh2^WT^* versus naïve *Ezh2^Y641F^* Treg cells. (G) Quantified frequencies of CD103^+^, PD1^+^, and TIGIT^+^ of YFP^+^RFP^+^ Treg cells stratified based on CD44 expression as a marker of Treg cell activation state from LN of Treg.*Ezh2^WT^/Ezh2^WT^* and Treg.*Ezh2^Y641^/Ezh2^WT^* female mosaic mice. (H-I) Gene set enrichment analysis of genes upregulated (H) or downregulated (I) in anti-CD3/anti-CD28 co-stimulated versus anti-CD3 stimulated CD4^+^ T cells (obtained from GSE39595) compared to CL2 segment-associated gene expression in naïve *Ezh2^Y641F^* Treg cells versus naïve *Ezh2^WT^* Treg cells (H) or compared to CL4 segment-associated gene expression in naïve *Ezh2^Y641F^* Treg cells versus naïve *Ezh2^WT^* Treg cells. Genomic data are obtained from three biological replicates. Flow cytometry date are mean ± SEM and representative of at least two independent experiments (n=8-11 mice total per genotype). Wilcoxon signed-rank test (E) or ordinary two-way ANOVA (G) was used. *p < 0.05, **p < 0.01, ***p < 0.001, and ****p<0.0001 (only statistically significant differences are described and non-significant data is not indicated). See also Figure S7.

### The redistribution of H3K27me3 in *Ezh2^Y641F^* Treg cells drives a gene expression pattern in naïve Treg cells that mimics CD28-activated *Ezh2^WT^* Treg cells

To better decipher how the specific changes in H3K27me3 modifications at genetic loci impacted the *Ezh2^Y641F^* Treg cell phenotype, we first generated a set of defined H3K27me3 domains based on H3K27me3 datasets from naïve, activated, *Ezh2^WT^*, or *Ezh2^Y641F^* Treg cells and identified unique, non-overlapping segments (Figure 7A and Figure S7A). We performed PCA using H3K27me3 enrichment based on all the unique segments identified (Figure 7B). This analysis revealed that PC2, which accounted for 26.6% of the variation, captured changes due to activation. Interestingly, naïve *Ezh2^Y641F^* Treg cells clustered more closely to activated *Ezh2^WT^* Treg cells than to naïve *Ezh2^WT^* Treg cells. The activated *Ezh2^Y641F^* Treg cells then further separated from all groups along PC2 (Figure 7B). Next, we scored the H3K27me3 signal for each segment and performed k-means clustering (k=4) of the comparison of all datasets to naive *Ezh2^WT^* Treg cells (Figure 7C). Cluster 1 (CL1) contained segments that gained H3K27me3 (red) in naïve *Ezh2^Y641F^*, activated *Ezh2^Y641F^,* and activated *Ezh2^WT^* Treg cells, whereas cluster 2 (CL2) contained segments that lost H3K27me3 (blue) in each group compared to naive *Ezh2^WT^* Treg cells. This showed that CL1 and CL2 contained segments that had changes in H3K27me3 levels in naive *Ezh2^Y641F^* Treg cells that mirrored H3K27me3 changes due to activation in *Ezh2^WT^* Treg cells. Thus, the expression of EZH2^Y641F^ in naïve Treg cells altered the H3K27me3 landscape to adopt similarities to that of activated Treg cells. This poising of naïve Tregs towards an activated Treg H3K27me3 phenotype in CL1 and CL2 may contribute to the enhanced activation and effector differentiation status observed in *Ezh2^Y641F^* Treg cells. Cluster 3 (CL3) contained segments that specifically lost H3K27me3 in both naïve and activated *Ezh2^Y641F^* Treg cells compared to naïve and activated *Ezh2^WT^* Treg cells, indicating that these segments were altered in response to the expression of EZH2^Y641F^, regardless of the Treg cells activation state. Cluster 4 (CL4) contained segments that gained H3K27me3 modifications specifically in naive *Ezh2^Y641F^* Treg cells, although these segments were largely erased in activated *Ezh2^Y641F^* compared to *Ezh2^WT^* Treg cells. PCA using H3K27me3 enrichment in CL1-and CL2-associated segments demonstrated that along PC1, which explains almost 50% of the variance, naïve *Ezh2^Y641F^* Treg cells cluster more closely to activated *Ezh2^WT^* Treg cells than to naïve *Ezh2^WT^* Treg cells (Figure S7B-C). Similar to our previous analysis (Figure 6D), for naïve *Ezh2^Y641F^* Treg cells, and to some extent for activated *Ezh2^Y641F^* Treg cells, we noticed a reduced fraction of domains overlapping with promoter regions, UTRs, and exons (Figure S7D). Furthermore, segments in CL1, which features a gain in H3K27me3 in naïve *Ezh2^Y641F^* Treg cells similar to activated Treg cells, showed a depletion of promoters, while segments in CL2, which features loss of H3K27me3 in naïve *Ezh2^Y641F^* Treg cells similar to activated *Ezh2^WF^* Treg cells, demonstrated an enrichment of promoters. This data again indicated that EZH2^Y641F^ activity promotes the loss of H3K27me3 at promoters and the gain of H3K27me3 elsewhere, which were changes normally associated with the Treg cell activated state.

To determine whether the H3K27me3 segment clusters correlated with changes in gene expression, we performed RNA-sequencing on naïve and activated *Ezh2^WT^* and *Ezh2^Y641F^* Treg cells. We then generated a “superset” of genes comprised of all the differentially expressed genes (DEGs) between any of the six pairwise comparisons between each condition (Figure 7D). Next, we identified genes from the superset whose promoters overlapped with each of the four H3K27me3 segment clusters (Figure 7C and S7E). Analysis of the expression of genes associated with the segments in each cluster revealed that the expression of genes associated with CL1 was decreased, whereas the expression of genes associated with segments in CL2 was increased in naïve *Ezh2^Y641F^* Treg cells compared to naïve *Ezh2^WT^* Treg cells (Figure 7E, left panel). Since CL1 and CL2 contained segments that increased or decreased in H3K27me3 enrichment with Treg activation (Figure 7C), these data indicated a clear anti-correlation between gene expression changes and changes in H3K27me3 enrichment in CL1 and CL2 with Treg cell activation (Figure 7E, right panel), which was in line with H3K27me3-mediated gene repression associated with each cluster. For the genes associated with CL3 and CL4, the most profound changes were observed in naïve and activated *Ezh2^Y641F^* Treg cells, with modestly increased expression of genes associated with CL3 and modest decreased expression of genes associated with CL4 (Figure 7E and S7F). Gene set enrichment analysis (GSEA) demonstrated that genes found to be downregulated in Treg cells compared to conventional CD4^+^ T cells from two publicly available datasets (GSE7852 and GSE7460) were enriched in genes that were downregulated in naïve *Ezh2^Y641F^* Treg cells, suggesting that *Ezh2^Y641F^* Treg cells expressed a stronger Treg cell phenotype compared to *Ezh2^WT^* Treg cells (Figure S7G). However, this positive correlation was not observed for the genes found to be upregulated in Treg cells compared to conventional CD4^+^ T cells, which is in line with the observation that EZH2 is essential for the establishment of gene repression at Foxp3-bound loci in Treg cells (Figure S7H) ^26,27^. Furthermore, a comparison of *Ezh2*-deficient Treg cells with *Ezh2^Y641F^* Treg cells, also demonstrated that genes found to have increased expression in either naïve or activated *Ezh2*-deficient compared to wild-type Treg cells, were enriched in genes with decreased expression in naïve *Ezh2^Y641F^* compared to naïve *Ezh2^WT^* Treg cells (Figure S7I). This confirmed an opposing effect in gene expression between Treg cells with reduced EZH2 activity versus increased EZH2 activity.

Analysis of the gene expression of the activation and effector differentiation markers identified by flow cytometry specifically in naïve *Ezh2^Y641F^* Treg cells revealed that in the naïve state, *Ezh2^Y641F^* Treg cells exhibited increased gene expression of *Itgae*, *Tigit*, *Pdcd1*, *Il10*, *Cxcr3*, *Ccr8*, *Ccr6*, and *Il1rl1* (but decreased expression of *Il2ra*) compared to naïve *Ezh2^WT^* Treg cells (Figure 7F). In line with this observation, we re-analyzed the surface expression of CD103, PD1, and TIGIT on flow samples that had been co-stained with CD44 to mark naïve (CD44^-^) versus activated (CD44^+^) *Ezh2^WT^* and *Ezh2^Y641F^* Treg cells from female Treg.*Ezh2^Y641F^/Ezh2^WT^* and Treg.*Ezh2^WT^/Ezh2^WT^* mice and found that each protein’s expression was increased both on naïve and activated *Ezh2^Y641F^* Treg cells compared to *Ezh2^WT^* Treg cells (Figure 7G). This data indicated that the expression of EZH2^Y641F^ drove Treg cells to adopt characteristics of activated Treg cells while still in a naïve state, and that once activated, EZH2^Y641F^ expression further enhanced Treg cell effector differentiation. We hypothesized that EZH2^Y641F^ expression poised naïve *Ezh2^Y641F^* Treg cells in a more activated state due to the increased activity of EZH2, which is normally induced by CD28 co-stimulation ^27^. GSEA comparing gene expression from naïve *Ezh2^Y641F^* versus *Ezh2^WT^* Treg cells to gene set signatures from T cells activated by anti-CD3/anti-CD28 (co-stimulation) compared to anti-CD3 alone (GSE39595) showed that genes from this dataset with increased expression due to co-stimulation were enriched in CL2-associated genes with increased expression in naïve *Ezh2^Y641F^* Treg cells (Figure 7H). Similarly, genes that were decreased in expression with co-stimulation were enriched in CL4-associated genes with decreased expression in naïve *Ezh2^Y641F^* Treg cells (Figure 7I). Comparison of global gene expression between naïve *Ezh2^Y641F^* and *Ezh2^WT^* Treg cells to the co-stimulation dataset also showed strong concordance with genes exhibiting decreased expression with co-stimulation, but not genes with increased expression, which was in line with EZH2’s role in the establishment of gene repression at Foxp3-bound loci in activated Treg cells (Figure S7J) ^26,27^. Overall, this suggests that in naïve *Ezh2^Y641F^* Treg cells, H3K27me3-associated genes take on gene expression patterns similar to CD28-co-stimulated Treg cells, thus poising naïve Treg cells to rapidly differentiate and migrate to organ tissues to function upon activation during an immune response.

## Discussion

Previous studies have demonstrated that Ezh2 deletion in Treg cells, which reduced H3K27me3 levels, impaired Treg cell function and caused autoimmunity ^27,30^. However, the direct impact of H3K27me3 levels on Treg cell function and whether increasing H3K27me3 deposition in Treg cells could enhance Treg cell function had not been examined. Here we generated Treg.*Ezh2^Y641F^* mice expressing an EZH2 SET-domain gain-of-function mutation (Y641F) specifically in Treg cells. We demonstrated that increased EZH2 function, and thus increased H3K27me3, increased Treg cell stability, promoted the generation of an effector differentiation Treg cell phenotype, and increased Treg cell migratory capacity to organ tissues. These characteristics allowed for a more rapid resolution of inflammation in the CNS upon acute induction of experimental autoimmune encephalomyelitis. Analysis of the genomic landscape of H2K27me3 modifications in naïve *Ezh2^Y641F^* Treg cells revealed that hyperactive EZH2 drove features of H3K27me3 modifications found in CD28-activated Treg cells. This is in line with previous work showing that CD28 co-stimulation induced EZH2 activity in Treg cells ^27^. Thus, hyperactive EZH2 initiates H3K27me3-mediated chromatin reorganization that poises naïve Treg cells in a state approaching that of activated Treg cells.

Mice with constitutively active EZH2 in Treg cells exhibited reduced frequencies of Treg cells compared to wild-type mice. Notably, this is in opposition to what was found in mice with deletion of *Ezh2* in Treg cells, where increased frequencies of Treg cells were observed ^27^. We hypothesize that these changes in the frequencies of Treg cells were indirect, resulting from compensatory feedback mechanisms to balance immune homeostasis. Therefore, the reduction in *Ezh2^Y641F/+^* Treg cells was the result of having more functional *Ezh2^Y641F/+^* Treg cells over a prolonged period of time, which could maintain immune homeostasis with fewer total Treg cells. In contrast, *Ezh2*-deficiency in Treg cells resulted in a greater demand for Treg cell output from the thymus to compensate for Treg cell functional insufficiencies ^27^. Interestingly, in the tamoxifen-inducible *Ezh2^Y641F^* activation model, such a decrease in Treg cells was not observed, and even an increase in Treg frequencies was detected in some experiments (data not shown). We hypothesize that at this timepoint, there is not enough time for the Treg cell compartment to contract in response to more functional *Ezh2^Y641F/+^* Treg cells. Such compensatory mechanisms may also explain why there was no difference in the course of EAE disease in mice constitutively expressing *Ezh2^Y641F^* Treg cells (data not shown), whereas mice with inducible expression of *Ezh2^Y641F^* in Treg cells led to a significantly improved reversal of EAE compared to mice with wild-type Treg cells (Figure 5A). In contrast, mice with *Ezh2*-deficient Treg cells showed the opposite phenotype in response to EAE, completely failing to resolve EAE ^27^. In addition, the frequency of MOG-specific Treg cells in the CNS was decreased in Treg-specific *Ezh2*-deficient mice, whereas the frequency of MOG-specific Treg cells in mice with *Ezh2^Y641F^* Treg cells was increased (Figure 5B). In both mice with *Ezh2^Y641F/+^* Treg cells and mice with *Ezh2*-deficient Treg cells, only recovery from EAE, but not EAE onset, was affected. Since *Ezh2*-deletion impacted Treg cells after activation and increased EZH2 function imparted increased effector differentiation in naïve Treg cells that promoted rapid localization to the CNS, EZH2 function in Treg cells appears to impact the resolution of inflammation in organ tissues rather than altering Treg cell capacity to control the priming of immune responses. Together, these data clearly indicate that EZH2 can act as an epigenetic switch by controlling the levels of H3K27me3 modifications in Treg cells, which with decreased H3K27me3, cause Treg cells to lose activity in organ tissues, whereas, with increased H3K27me3, Treg cells gain tissue homing and maintenance in organ tissues.

Increased EZH2 activity in *Ezh2^Y641F/+^* Treg cells led to the increased expression of the homing receptors CXCR3, CCR8, CCR6, and IL33R, thereby promoting Treg cell migration to organ tissues. This was revealed in two different competitive mouse models wherein WT and *Ezh2^Y641F/+^* Treg cells co-populated mice and *Ezh2^Y641F/+^* Treg cells overrepresented the Treg cell pool in organ tissues and sites of inflammation. Notably, we found the expression of each receptor to be expressed on different Treg cells, generating a heterogeneous population of effector differentiated Treg cells. Treg cells can be divided into multiple distinct subsets with unique migratory and functional characteristics ^90^. For example, expression of CXCR3 on Treg cells supports Treg cell migration to sites of Th1 inflammation, while Treg control of Th2 inflammatory responses requires the expression of CCR4 and CCR8 ^71–74^. Alternatively, at sites of Th17 inflammation, recruited Treg cells have been shown to express CCR6 ^70^. Besides direct control of T cell responses, Treg cells are also involved in tissue protection and repair, in particular via the production of amphiregulin in response to the alarmin IL-33 ^75,78^. Thus, the presence of IL-33R^+^ (ST2^+^) expressing Treg cells has been shown to be critical for wound healing and repair after tissue damage ^75–78^. Therefore, it appears that *Ezh2^Y641F/+^* Treg cells are poised to differentiate into a heterogeneous population of effector Treg cells endowed with features to promote migration and retention in organ tissues. Data from this study is consistent with these effects being due to changes in the H3K27me3 landscape induced by EZH2^Y641F^. However, extranuclear EZH2 activities have also been described that control actin polymerization to regulate cell adhesion and migration, which may have also contributed to the enhanced migration phenotype of *Ezh2^Y641F/+^* Treg cells ^91,92^.

EZH2 has been demonstrated to play a key role in collaboration with FOXP3 in maintaining the signature Treg cell transcriptome and promoting Treg cell function after activation ^26–28^. Analysis of the H3K27me3 landscape of Treg cells containing increased EZH2 function revealed a global redistribution of H3K27me3 away from the TSS and PRC2 nucleation sites towards intergenic regions. Promoter regions where H3K27me3 levels were depleted in *Ezh2^Y641F/+^* Treg cells were mostly associated with embryonic and neuronal gene sets, which is in agreement with the important role of PRC2 during neural development ^93,94^. Homeobox genes also significantly lost H3K27me3 levels in *Ezh2^Y641F/+^* Treg cells compared to *Ezh2^WT^* Treg cell (data not shown). More in-depth analysis of H3K27me3 segments present within naïve and activated *Ezh2^Y641F^* Treg cells compared to *Ezh2^WT^* Treg cells demonstrated that two clusters of H3K27me3 segments present in activated Treg cells were already present in naïve *Ezh2^Y641F^* Treg cells, suggesting that increased activity of EZH2 poises Treg cells to adopt an activated phenotype. The global re-distribution of H3K27me3 away from the promoter/TSS towards intergenic regions that we observed in *Ezh2^Y641F^* Treg cells was also described in developing B cells and embryonic stem cells (ESCs) expressing *Ezh2^Y641F^* ^43,85^. Interestingly, *Ezh2^Y641F^* expression enhanced B cell differentiation, and the H3K27me3 profile of ESCs expressing EZH2^Y641F^ resembled that of ESCs undergoing differentiation, which is in line with our observations that *Ezh2^Y641F^* Treg cells appeared more differentiated than *Ezh2^WT^* Treg cells. However, phenotypic effects of *Ezh2^Y641F^* expression seem to be dependent on the cellular developmental stage, as expression of EZH2^Y641F^ in developed mature B cells *in vitro* led to increased H3K27me3 modifications at promoter areas and an irreversible block in differentiation ^40^. In the constitutive *Ezh2^Y641F^* model as well as the inducible *Ezh2^Y641F^* model used in this study, hyperactive *Ezh2^Y641F^* expression in Treg cells was induced only after Foxp3 expression and Treg cell differentiation; thus, EZH2 activity was increased only in established Treg cells. This eliminated any confounding effects of hyperactive *Ezh2^Y641F^* expression in Treg precursor cells during T cell development in the thymus. Pluripotent ESCs have lower H3K27me3 levels compared to more differentiated cells, and it has been suggested that the length of the G1 phase of the cell cycle, which is short in ESCs, influences the distribution and enrichment of H3K27me3 genome-wide ^95–98^. T cell stimulation leads to an acceleration in cell cycle progression, with T cells dividing every 8-10 hours during the peak of a response ^99,100^. Therefore, EZH2 expression increases to maintain H3K27me3 levels during proliferation, which acts to maintain cell state identity ^101^. Here we show that increased EZH2 activity with the Y641F mutation preemptively mimics many of the changes in the H3K27me3 landscape that occur with Treg cell activation, thereby poising Treg cells with an activated effector differentiated phenotype while still in a naïve, pre-activated state.

The data presented in this study suggests the potential for drugs that increase H3K27me3 levels in Treg cells to be therapeutic in the settings of autoimmunity or transplantation tolerance. One such compound, GSK-J4, which inhibits the H3K27 demethylase KDM6B/JMJD3, has already been shown to reduce the severity of EAE and ameliorate DSS-induced acute colitis in mice ^102–104^. Although GSK-J4 treatment favored Treg cell differentiation, stability, and suppressive function *in vitro*, this effect was only visible in the presence of dendritic cells (DCs), which has led to the hypothesis that GSK-J4 activity is due to driving tolerogenic DCs that then promote Treg cell function, rather than GSK-J4 acting directly on the Treg cells themselves ^102,103^. Analysis of *Jmjd3*-deficient T cells demonstrated that *in vitro* Treg cell differentiation was impaired ^105^. However, *Jmjd3*-deficient natural Treg cells were equally effective as WT Treg cells in an *in vivo* colitis model ^105^. Furthermore, mice with *Jmjd3*-deficient Treg cells exhibited more rapid tumor outgrowth, which suggests that *Jmjd3*-deficient Treg cells were more suppressive than wild-type Treg cells in the setting of cancer ^30^. Nevertheless, it is complicated to make direct comparisons between the effect of *Jmjd3*-deficiency and increased EZH2 activity on Treg cell behaviors since JMJD3 and EZH2 act in distinct protein complexes and may have distinct genetic targets ^106,107^. Thus, our use of a hyperactive *Ezh2^Y641F^* allele here, likely served as a better test of the role of increased EZH2 function in Treg cells. However, non-canonical protein-protein interactions of the hyperactive EZH2^Y641F^ protein can cause it to act as a transcriptional activator in certain contexts, making the gain of functional activities a potential driver of the phenotypes of *Ezh2^Y641F^* Treg cells^108^. Ultimately, a deeper exploration of JMJD3 inhibition in Treg cells is warranted and will determine whether this actionable drug target could enhance Treg cell suppression for the treatment of autoimmunity or transplantation tolerance.

Taken together, we have demonstrated that increased EZH2 function in Treg cells promotes their capacity to suppress autoimmunity by poising Treg cells for rapid effector differentiation and migration to organ tissues during an immune response. Increased EZH2 function induced an H3K27me3 landscape that mimicked features of CD28-activated Treg cells, indicating that EZH2 activity can directly promote aspects of Treg cell activation in the absence of extracellular stimulating cues. Therefore, drugs that can increase H3K27me3 levels in Treg cells could be a promising therapeutic approach to promote immune tolerance in the settings of transplantation or autoimmunity.

## Acknowledgements

We thank Djem Kissiov and David Raulet for sharing CUT&RUN protocols and reagents. We thank Genevia Technology, specifically Grigorios Georgolopoulos, for assistance in analyzing the H3K27me3 CUT&RUN data. We also thank Hector Nolla, Alma Valleros and Kartoosh Heydari of the UC Berkeley Cancer Research Laboratory Flow Cytometry Facility and the Functional Genomics Laboratory of UC Berkeley. Furthermore, we thank all the members of the DuPage Lab for providing feedback on the research approach and critically reviewing the manuscript. This research was supported by a ZonMw Rubicon fellowship #45219210 (to J.G.C.P), National Institute of Health grants 1DP2CA247830-01 (to M.D.) and R35GM133434 (to S.R.), American Cancer Society grant RSG-22-026-01 (S.R.). and the RNA Bioscience Initiative, the University of Colorado School of Medicine. S.R. is a Pew-Stewart Scholar for Cancer Research, supported by the Pew Charitable Trusts and the Alexander and Margaret Stewart Trust. M.D. is a Pew-Stewart Scholar and a St. Baldrick’s Scholar with generous support from Hope with Hazel.

## Author contributions

Conceptualization and methodology, M.D. and J.G.C.P.; investigation, J.G.C.P, S.S., M.O., S.R., and M.D.; software: S.R.; writing – original draft, J.G.C.P and M.D; writing – review & editing, J.G.C.P., S.R., M.D.; supervision, J.G.C.P., S.R., and M.D.; funding acquisition, M.D. and S.R. The authors declare no competing interests.

## Resource availability

### Lead contact

Further information and requests for resources and reagents should be directed to and will be fulfilled by the lead contact, Michel DuPage.

### Materials availability

*Foxp3-GFP-hCre;Ezh2^Y641F/+^*, *Foxp3^GFP-DTR^/Foxp3^YFP-Cre^*; *Ezh2^Y641F/+^*, and *Foxp3^eGFP-Cre-ERT2^*; *Ezh2^Y641F/+^* mice were generated for this study and will be made available from the lead contact upon request.

## Methods

### Mice

All mice used were bred onto a C57BL/6 background a minimum of ten generations. All mouse experiments used comparisons between littermates or age-matched control mice. *Ezh2^Y641F/+^* mice were generated and provided by Dr. Sharpless (University of North Carolina School of Medicine) and crossed to *Foxp3-Cre* driver alleles ^43^. *Foxp3-GFP-hCre* were kindly provided by Dr. Bluestone (University of California, San Francisco; JAX:023161) ^45^. *Foxp3^GFP-DTR^, Foxp3^YFP-Cre^*, and *Foxp3^eGFP-Cre-ERT2^* mice were a gift from Dr. Rudensky (Memorial Sloan Kettering Institute; JAX:016958, JAX:016959, and JAX:016961, respectively) ^79,80,109,110^. All the experiments were conducted according to the Institutional Animal Care and Use Committee guidelines of the University of California, Berkeley.

### Cell isolation and flow cytometry

Single cell suspensions from lymphoid organs were prepared by mechanical disruption in ice-cold PBS buffer containing 2% FCS and passing them through 40 μm filters. In addition, spleens were subjected to red blood cell lysis using ACK buffer (150mM NH4Cl, 10mM KHCO3, 0.1mM Na2EDTA, pH7.3). Isolation of lymphocytes from the spinal cord and cerebellum (CNS) of mice with EAE was done as described previously. In brief, after left ventricle perfusion spinal cords were extruded by flushing the vertebral canal with PBS and cerebella were removed. Spinal cords and hind cerebella were diced and incubated in Hank’s balanced salt solution (HBSS) containing 25 mM HEPES (Thermofisher Scientific), 320 U/mL Collagenase D (Roche), and 50 ug/mL DNase I (Roche) for 30 min, 37 °C, while shaking. Homogenates were resuspended in 30% isotonic Percoll (VWR), underlaid with 70% Percoll, and centrifuged at 2400 rpm at RT, 30 min. Mononuclear cells were collected from the Percoll interphase, washed twice HBSS containing 2% FCS and incubated with Human Trustain Fcx (Biolegend) for 20 min on ice. CNS-infiltrating cells were stained with MOG38–49-I-Ab-APC tetramer (NIH Tetramer Core) for 2h at RT, followed by staining with other antibodies on ice. Dead cells were stained with Live/Dead Fixable Violet or Aqua Dead Cell Stain kit (Molecular Probes) in PBS for 20 minutes at 4°C. Cell surface antigens were for 30 min at 4°C using a mixture of fluorophore-conjugated antibodies. Surface marker stains for murine samples were carried out with anti-mouse CD45 (30-F11, BioLegend), anti-mouse CD4 (RM4-5, BioLegend), anti-mouse CD8a (53-6.7, BioLegend), anti-mouse PD1 (RMP1-30, BioLegend), anti-mouse TIGIT (Vstm3, Biolegend), anti-mouse CD69 (H1.2F3, Biolegend), anti-CD25 (PC61, BioLegend), anti-mouse CD103 (2E7, BioLegend), anti-mouse CXCR3 (CXCR3-173, BioLegend), anti-mouse CCR8 (SA214G2), anti-mouse CCR6 (29-2L17, Biolegend), anti-mouse IL33R (ST2) (RMST2-2, eBioscience) in PBS 2% FCS. Prior to intracellular staining, cells were either fixed using the Foxp3/Transcription Factor Staining Buffer Kit (Tonbo) or using 4% PFA, to preserve fluorescent reporter expression, followed by treatment with 0.1% Triton to permeabilize the cells. Intracellular staining was performed using anti-mouse Foxp3 (FJK-16S, eBioscience), anti-histone H3 (D1H2, Cell Signaling Technology), anti-tri-methyl-histone H3 (Lys27) (C36B11, Cell Signaling Technology), anti-mouse TNF-a (MP6-XT22, BioLegend), anti-mouse IFNg (XMG1.2, eBioscience), anti-IL10 (JES5-16E3, Biolegend) for 1h, at 4°C, according to manufacturer’s instructions. Cytokine staining was performed with 2 x 10^6^ cells after 3.5h of *in vitro* stimulation in Opti-MEM media supplemented with Golgiplug (BD Biosciences), 10 ng/ml phorbol 12-myristate 13-acetate (PMA) (Sigma), and 0.25 μM ionomycin (Sigma). Flow cytometry was performed on an BD LSR Fortessa X20 (BD Biosciences) or LSRFortessa (BD Biosciences) and datasets were analyzed using FlowJo software (Tree Star).

### Tamoxifen treatment

Tamoxifen (Sigma-Aldrich) was resuspended at a concentration of 40 mg/mL in corn oil (Fisher Science Education) and heated for 30 min at 65°C to achieve a completely dissolved solution. *Foxp3^eGFP-Cre-ERT^;Ezh2^+/+^* and *Foxp3^eGFP-Cre-ERT^;Ezh2^Y641F/+^* mice were treated twice, 2 or 3 days apart, with 4 mg tamoxifen via oral gavage and 14 days after the 2^nd^ treatment lymphoid tissues were harvested and cells were analyzed and/or isolated for experimental purposes.

### Experimental Autoimmune Encephalomyelitis model

Fourteen days after tamoxifen treatment, mice were immunized subcutaneously with 100 μl of emulsified Complete Freund adjuvant (BD Difco) supplemented with 4 mg/ml Mycobaterium tuberculosis H37Ra (BD Difco) and 200 μg MOG35-55 peptide (MEVGWYRSPFSRVVHLYRNGK, Genemed Synthesis) and received intraperitoneal injections of 200 ng Pertussis Toxin from *Bordetella pertussis* (List biological Laboratories) at the time of immunization and 48h later. Clinical disease was assessed by the scoring of ascending hind-limb paralysis as follows: no signs, score 0; paralysis of tail, score 1; hind-limb weakness, score 2; paralysis of one hind-limb, score 3; paralysis of both hind-limb, score 4; and moribund mouse, score 5.

### In vitro suppression assay

Spleens and lymph nodes were collected from *Foxp3^eGFP-Cre-ERT^;Ezh2^+/+^* and *Foxp3^eGFP-Cre-ERT^;Ezh2^Y641F/+^* mice treated with tamoxifen 14 days earlier. Single cell suspensions were generated and enriched for CD4^+^ T cells by negative selection using EasySep magnetic bead kit (STEMCELL Technologies) and stained with anti-mouse CD4 (RM4-5, BioLegend), anti-CD25 (PC61, BioLegend), anti-CD62L (MEL1-14, Biolegend), anti-CD357 (GITR) (DTA-1, Biolegend). Treg cells (CD4^+^Foxp3^GFP+^CD25^+^GITR^+^) and naïve CD4^+^ T cells (CD4^+^CD62L^-^ cells) were sorted using an Aria Fusion sorter (BD Biosciences) with a 70μm nozzle. Naïve CD4^+^ T cells were labeled with CellTrace Far Red Cell Proliferation Kit (Thermo Fisher Scientific) and cultured with different ratios of Treg cells and anti-CD90.2 (30-H12, Biolegend)-depleted splenocytes that were pre-incubated for 20 min with 1 mg/ml anti-CD3 antibody in DMEM medium supplemented with 10% FBS (Hyclone), 1% non-essential amino acids, 1 mM sodium pyruvate, 2 mM L-glutamine, 10 uM HEPES and 55 μM β-ME. The percentage of undivided cells (no dilution of CellTrace Far Red) was analyzed 3 days later.

### Adoptive transfer experiments

Spleens and lymph nodes were collected from *Foxp3-GFP-hCre;Ezh2^+/+^* and *Foxp3-GFP-hCre;Ezh2^Y641F/+^* mice and Treg cells (CD4^+^Foxp3^GFP+^R26^RFP+^CD62L^+^) were sorted as described above. Treg cells were activated with anti-CD3 and anti-CD28 coated beads (Invitrogen) at a ratio of 1:3 (cell:bead) in the presence of 2000 IU/mL recombinant human IL-2 (TECIMTM, Hoffman-La Roche provided by NCI repository, Frederick National Laboratory for Cancer Research) for 7 days in DMEM medium supplemented as described and kept at a concentration of 10^6^ cells/ml. Treg cells were stained with 5 uM ViaFluor 405 (Biotum) or 1uM CellTrace Far Red (Molecular Probes) dyes according to the manufacturer’s protocol, but including 5% FBS. Dyes were interchanged between experiments to prevent bias in the results due to potential differences in staining. Treg cells were co-transferred to WT mice at 1:1 Ezh2^WT^ to Ezh2^Y641F^ ratio by intravenous injection. 24 hours later, mice were intravenously injected with 0.2 ug CD45-BV785 and euthanized 5 minutes later after which tissues were removed for flow cytometry analysis.

### CUT&RUN

CUT&RUN was performed as described^111^. Briefly, 350,000 – 500,000 Treg cells (sorted as CD4^+^Foxp3^GFP+^RFP^+^CD62L^+^ from *Foxp3-GFP-hCre;Ezh2^Y641F/+^;R26^RFP^* or *Foxp3-GFP-hCre;Ezh2^+/+^;R26^RFP^* mice) or 500.000 CD4^+^ T effector cells (sorted as CD4^+^Foxp3^GFP-^RFP^-^CD62L^+^) were washed and immobilized on Con A beads (Bangs Laboratories) and permeabilized with wash buffer containing 0.01% w/v Digitonin (Sigma-Aldrich) either directly after sorting or after 4 days of activation with anti-CD3 and anti-CD28 coated beads at a ratio of 1:3 (cell:bead) in the presence of 200 IU/mL (CD4^+^ T effector cells) or 2000 IU/mL (Treg cells) recombinant human IL-2 in DMEM medium supplemented as described. Cells were incubated rotating for 2 hr at 4°C with 1 uL anti-tri-methyl-histone H3 (Lys27) (C36B11, Cell Signaling Technology) or normal rabbit IgG (#2729, Cell Signaling Technologies). Permeabilized cells were washed and incubated with pA-MNase (kindly provided by the Henikoff lab) at a concentration of 700 ng/mL for 10 min at room temperature, while rotating. After washing, cells were incubated at 0°C and MNase digestion was initiated by adding 2 mM CaCl^2^. After 30 min, the reaction was stopped by the addition of EDTA and EGTA and 2 pg/mL DNA from *Saccharomyces cerevisiae* micrococcal nuclease-treated chromatin (kindly provided by the Henikoff lab) was added ad spike-in DNA for calibration. Chromatin fragments were released by incubation at 37°C for 10 min, purified by overnight proteinase K digestion at a concentration of 150 μg/mL with 0.1% wt/vol SDS at 55°C. DNA was purified by phenol/chloroform extraction followed by PEG-8000 precipitation (final concentration of 20% wt/vol) using Sera-mag SpeedBeads (Fisher). Libraries were prepared using the NEBNext Ultra II DNA library prep kit for Illumina (New England Biolabs) according to manufacturer’s instructions with the following specifications and modifications. The entire preparation of purified CUT&RUN fragments from a reaction were used to create libraries. NEBNext adaptor for Illumina from NEBNext Multiplex Oligos for Illumina (New England Biolabs) was diluted 25-fold in TBS buffer. Size selection was performed with AmpureXP beads (Agencourt), adding 0.4 X volumes to remove large fragments, after which supernatant was recovered and 0.6 X volumes of AmpureXP beads and 0.6 X volumes of PEG-8000 (20% wt/vol PEG-8000, 2.5 M NaCl) were added for recovery of smaller fragments. Adapter-ligated libraries were amplified for 15 cycles using NEBNext Ultra II Q5 Master Mix using the universal primer and an indexing primer provided with the NEBNext Multiplex Oligos for Illumina. Amplified libraries were further purified with the addition of 1.1 X volumes of AmpureXP beads to remove adapter dimer and eluted in 25 μL H^2^O. Libraries were quantified by Qubit (ThermoFisher) and Bioanalyzer (Agilent) and sequenced 150 bp paired-end on an Illumina NovaSeq 6000 by Genewiz (Azanta Life Sciences).

### CUT&RUN analysis

For analysis in Figure 6, samples were aligned to GRCm38 (mm10) using *bowtie2* in *local* mode with the following parameters: -I 10 -X 700 –phred33 -very-sensitive-local and also aligned to spike-in genome (SacCer R64) using bowtie2 with the following parameters: -dovetail -phred33^112^. Improperly mapped reads and mates were discarded, and duplicates were marked with Picard. Library complexity was calculated according to ENCODE DCC guidelines. Coverage was scaled based on spike-in ratios (scaling factor: divide spike-in reads in each sample by the lowest number of spike-in reads). Peaks were called using epic2 considering all replicates per sample and providing the corresponding IgG input as control ^113,114^. The option –keep-duplicates was set to include reads marked as duplicates by Picard. Peaks were filtered by keeping peaks with peaks with Benjamini-Hochberg corrected p-value (q-value) < 1e-20. Pairwise overlaps between samples were calculated using BEDOPS and considered as overlaps all the peaks where at least 20% of the peak in the reference set overlaps with a peak in the map set ^115^. Peaks were annotated with HOMER using the annotatePeaks.pl command. To calculate genome-wide differences in H3K27me3, the genome was segmented to 1kb bins and the log2 ratio of H3K27me3 over IgG was quantified for every bin. To identify genes with differential H3K27me3 levels, the log2 ratio of each sample over IgG H3K27me3 signal was used and the average ratio over each gene body or gene promoter was calculated for each gene (55,398 genes). For every pairwise comparison, the group means were tested using the Wald test to simulate the analysis performed by DESeq2 ^116^. P-values were adjusted using the Benjamini-Hochberg FDR correction. Results were filtered to include genes with adjusted p-value < 0.05, and absolute log2FoldChange (difference between the log2 ratio of the test group and the reference group) > 0.58.

For Figure 7, CUT&RUN sequencing reads were first trimmed using Cutadapt: Illumina adapter sequences were removed and reads trimmed to 140 bp ^117^. Reads less than 35 bp were discarded. Trimmed reads were aligned to the mm10 version of *Mus musculus* genome using bowtie2. Samtools and bedtools were used to process aligned reads from sam to bed files ^118,119^. Duplicate reads were discarded for further analysis if the reads had the same start and end coordinates. Coverage at 100 bp windows genome-wide was calculated as the number of reads that mapped at that window, normalized by the factor N:

N = 10,000/(Total number of spike-in reads)

10,000 was a number chosen arbitrarily. The spike-in normalized read counts were then smoothed with a running average spanning +/- 1000 bp around each 100 bp bin. The distribution of normalized read counts in 100 bp windows genome-wide was generated, and a “domain cutoff” was determined as the normalized read count that is greater than the normalized read count of 95% of the windows. Domains were called by linking adjacent windows with a normalized read count ≥ domain cutoff. To account for short disruptions due to mappability issues, jumps of up to 750 bp were allowed while linking windows. A log2 ratio of H3K27me3 enrichment over IgG enrichment was calculated for all the putative domains. Those domains with a log2 enrichment greater than 2 (4-fold enrichment over IgG) were used for all downstream analyses. For performing disjoin and reduce operations, the GenomicRanges package in R was used ^120^. For chromatin accessibility analysis, H3K27me3 segments with a ≥ 2-fold change in H3K27me3 between naïve *Ezh2^Y641F^* and *Ezh2^WT^* Treg cells were identified and divided into four quartiles. Normalized ATAC-seq read counts were plotted for each quartile and the frequency of H3K27me3 segments overlapping with ATAC-seq peaks separated by each quartile was calculated.

### RNA-sequencing

RNA was extracted using the RNAeasy Micro kit (Qiagen) according to manufacturer’s instructions from Treg cells (sorted as CD4^+^Foxp3^GFP+^RFP^+^CD62L^+^ from *Foxp3-GFP-hCre;Ezh2^Y641F/+^;R26^RFP^* or *Foxp3-GFP-hCre;Ezh2^+/+^;R26^RFP^* mice) directly after sorting or after 4 days of activation with anti-CD3 and anti-CD28 coated beads (Invitrogen) at a ratio of 1:3 (cell:bead) in the presence of 2000 IU/mL recombinant human IL-2 (TECIMTM, Hoffman-La Roche provided by NCI repository, Frederick National Laboratory for Cancer Research) in DMEM medium supplemented as described. Libraries were prepared using KAPA mRNA HyperPrep Kit according to manufacturer’s instructions with the following specifications. 120 ng RNA was used as starting material, mRNA fragmentation was performed for 7 minutes at 94 °C, 1.5 μM xGEN UDI-UMI adapters (Integrated DNA Technologies) were used for adapter ligation, and adapter-ligated libraries were amplified for 13 cycles. Libraries were quantified by Qubit (ThermoFisher) and Bioanalyzer (Agilent) and sequenced 100 bp single-end on an Illumina NovaSeq 6000.

### RNA-seq analysis

Transcripts were quantified using Salmon (v1.9.0) with transcript definitions from ENSEMBL (release 102, Mus_musculus.GRCm38.cdna.all.fa.gz) ^121^. Salmon quantification was used directly in DESeq2 with four groups (three replicates each): naïve *Ezh2^WT^*, naïve *Ezh2^Y641F^*, activated *Ezh2^WT^*, and *Ezh2^Y641F^* Treg cells ^116^. We then performed all six possible pairwise comparisons between the four groups to identify genes with significant log2 foldchange (adjusted p-value < 0.05). This superset of significantly changing genes was used in downstream analyses. For plotting gene quantifications, we transformed the count data using rlog (regularized logarithm) function in DESeq2 with the setting “blind=FALSE”.

### Over-representation analysis and Gene set enrichment Analysis

Over-representation analysis was performed using WebGestalt ^122^. Gene set enrichment analysis was performed using GSEA ^123^. Significance of the enrichment was calculated based on 1000 cycles of permutations and the normalized enrichment score and p-value are annotated. Gene sets used to perform enrichment analysis are specified in the figure legend.

### Statistical Methods

p values were obtained from unpaired two-tailed Student’s t tests for all statistical comparisons between two groups, and data were displayed as mean ± SEM. For EAE disease score over time, a two-way repeated measured ANOVA was used. For comparisons between more than two groups, ordinary one-way ANOVA was used. Significance of pairwise comparisons of H3K27me3 signal in gene bodies and promoters was tested using a Wald test. Wilcoxon-signed rank tests were used to test if the distribution of fold changes in gene expression were significantly different shifted compared to 0. Over- and underrepresentation analysis was performed using hypergeometric test after multiple testing correction. p values are denoted in figures by *p < 0.05, **p < 0.01, ***p < 0.001, and ****p < 0.0001.

## Supplemental Figure Legends

**Figure S1. Expression of *Ezh2^Y641F^* in Treg cells increases H3K27me3 levels in the spleen.**

(A-B) Normalized H3K27me3/Histone H3 levels of CD4^+^Foxp3^+^ (A) or CD4^+^Foxp3^-^ (Teff) (B) cells in spleen of Treg.*Ezh2^WT^* and Treg.*Ezh2^Y641F^* mice.

(A) (C) Frequency of CD4^+^GFP^+^RFP^+^ cells of CD4^+^ and LIVE cells in spleen of Treg.*Ezh2^WT^* and Treg.*Ezh2^Y641F^* mice.

(B) (D) Frequency of CD8^+^, CD8^+^CD44^+^, CD4^+^ Teff, and CD4^+^CD44^+^ Teff in spleen of Treg.*Ezh2^WT^* and Treg.*Ezh2^Y641F^* mice.

(C) (E) Representative flow cytometry plots and quantified frequencies of TNFα^+^IFNγ^+^ of CD8^+^ and CD4^+^ Teff cells in the spleen of Treg.*Ezh2^WT^* and Treg.*Ezh2^Y641F^* mice with (+) or without (-) *in vitro* stimulation with PMA and ionomycin.

Data are mean ± SEM and representative of at least two independent experiments (n=12-13 mice total per genotype). Unpaired two-tailed Student’s t-test; p < 0.05, **p < 0.01, ***p < 0.001, and ****p<0.0001 (only statistically significant differences are noted and non-significant data is not indicated).

**Figure S2. *Ezh2^Y641F^* Treg cells exhibit increased activation and effector differentiation in the spleen.**

(A) Representative flow cytometry plots and quantified frequencies of IL-10^+^ Treg cells from spleen of Treg.*Ezh2^WT^* and Treg.*Ezh2^Y641F^* mice.

(B) Representative flow cytometry plots and quantified frequencies of TIGIT^+^ Treg cells in spleen of Treg.*Ezh2^WT^* and Treg.*Ezh2^Y641F^* mice.

(C) Quantified frequencies of PD1^+^, CD69^+^ and CD25^+^ Treg cells in spleen of Treg.*Ezh2^WT^* and Treg.*Ezh2^Y641F^* mice.

(D) Representative flow cytometry plots and quantified frequencies of CD103^+^ of Treg cells in spleen of Treg.*Ezh2^WT^* and Treg.*Ezh2^Y641F^* mice.

(E) UMAP analysis demonstrating the expression of CD103, TIGIT, PD1, CD69, and CD25 in Treg cells from LN of Treg.*Ezh2^WT^* mice.

(F) Representative flow cytometry plots and quantified frequencies of CXCR3^+^ in spleen of Treg.*Ezh2^WT^* and Treg.*Ezh2^Y641F^* mice.

(G) Quantified frequencies of CCR8^+^ and IL33R^+^ Treg cells in spleen of Treg.*Ezh2^WT^* and Treg.*Ezh2^Y641F^* mice.

(H) UMAP analysis demonstrating the expression of CXCR3, CCR8, CCR6, and IL33R in Treg cells from LN of Treg.*Ezh2^WF^* mice.

Data are mean ± SEM and representative of at least two independent experiments (n=12-13 mice total per genotype). Unpaired two-tailed Student’s t-test; *p < 0.05, **p < 0.01, ***p < 0.001, and ****p<0.0001 (only statistically significant differences are noted and non-significant data is not indicated).

**Figure S3. *Ezh2^Y641F^* Treg cells have a competitive advantage over WT Treg cells in the spleen.**

(A) Normalized ratio YFP^+^RFP^+^/GFP^+^RFP^-^ Treg cells in the spleen of Treg.*Ezh2^WT^/Ezh2^WT^* and Treg.*Ezh2^Y641F^/Ezh2^WT^* female mosaic mice.

(B) Normalized H3K27me3/Histone H3 levels of CD4^+^Foxp3^+^ Treg cells in the spleen of Treg.*Ezh2^WT^/Ezh2^WT^* compared to Treg.*Ezh2^Y641^/Ezh2^WT^* mice.

(C) Frequency of CD4^+^Foxp3^+^ cells in spleen of Treg.*Ezh2^WT^/Ezh2^WT^* and Treg.*Ezh2^Y641F^/Ezh2^WT^* mice.

(D) Quantified frequencies of total CD8^+^ T cells, CD44^+^CD8^+^ T cells, and TNFα^+^IFNγ^+^CD8^+^ T cells (with PMA/Iono restimulation) (top row) and total CD4^+^ Teff cells, CD44^+^CD4^+^ Teff cells, and TNFα^+^IFNγ^+^CD4^+^ Teff cells (with PMA/Iono restimulation) (bottom row) in spleen of Treg.*Ezh2^WT^/Ezh2^WT^* and Treg.*Ezh2^Y641F^/Ezh2^WT^* mice.

(E) Frequency of YFP^+^RFP^+^ Treg cells (as a percentage of CD4^+^RFP^+^ cells) in spleen of Treg.*Ezh2^WT^/Ezh2^WT^* and Treg.*Ezh2^Y641F^/Ezh2^WT^* mice.

(F) Quantified frequencies of TIGIT^+^ and PD1^+^ of YFP^+^RFP^+^ Treg cells in the spleen of Treg.*Ezh2^WT^/Ezh2^WT^* and Treg.*Ezh2^Y641F^/Ezh2^WT^* mice.

(G) Quantified frequencies of CD103^+^, CD69^+^, CD25^+^ of YFP^+^RFP^+^ Treg cells in the spleen of Treg.*Ezh2^WT^/Ezh2^WT^* and Treg.*Ezh2^Y641F^/Ezh2^WT^* mice.

(H) Quantified frequencies of CXCR3^+^, CCR8^+^, CCR6^+^, and IL33R^+^ of YFP^+^RFP^+^ Treg cells in the spleen of Treg.*Ezh2^WT^/Ezh2^WT^* and Treg.*Ezh2^Y641F^/Ezh2^WT^* mice.

Data are mean ± SEM and representative of at least two independent experiments (n=8-11 mice total per genotype). Unpaired two-tailed Student’s t-test; *p < 0.05, **p < 0.01, ***p < 0.001, and ****p<0.0001 (only statistically significant differences are noted and non-significant data is not indicated).

**Figure S4. Acute induction of *Ezh2^Y641F^* does not immediately impact T effector cell homeostasis.**

(A) Normalized H3K27me3/Histone H3 levels of CD4^+^Foxp3^+^ Treg cells in the spleen of *Ezh2^Y641F^*-inducible Treg.*iEzh2^WT^* compared to Treg.*iEzh2^Y641F^* mice two weeks after tamoxifen treatment.

(B) Quantified frequencies of CD4^+^Foxp3^+^ Treg cells in the spleen of *Ezh2^Y641F^*-inducible Treg.*iEzh2^WT^* and Treg.*iEzh2^Y641F^* mice two weeks after tamoxifen treatment.

(C) Frequencies of undivided CD4^+^ T effector cells after co-culture with indicated ratio of Treg/Teff cells (-indicates no Treg cells) (one of three biological replicates shown). Treg cells were FACS purified from *Ezh2^Y641F^*-inducible Treg.*iEzh2^WT^* and Treg.*iEzh2^Y641F^* mice two weeks after tamoxifen treatment.

(D) Quantified frequencies of total CD8^+^ T cells, CD44^+^CD8^+^ T cells, and TNFα^+^IFNγ^+^CD8^+^ T cells (with PMA/Iono restimulation) (top row) and total CD4^+^ Teff cells, CD44^+^CD4^+^ Teff cells, and TNFα^+^IFNγ^+^CD4^+^ Teff cells (with PMA/Iono restimulation) (bottom row) in spleen of Treg.*iEzh2^WT^* and Treg.*iEzh2^Y641F^* mice two weeks after tamoxifen treatment.

Data are mean ± SEM and representative of at least two independent experiments (n=10-14 mice total per genotype), unless noted otherwise. Unpaired two-tailed Student’s t-test; *p < 0.05, **p < 0.01, ***p < 0.001, and ****p<0.0001 (only statistically significant differences are noted and non-significant data is not indicated).

**Figure S5. Enhanced effector differentiation of *Ezh2^Y641F^* in LN is associated with increased migration to organ tissues.**

(A) Quantified frequencies of CD103^+^, TIGIT^+^, and CD69^+^ Treg cells within the draining LN (top row) or CNS tissues (bottom row) of Treg.*iEzh2^WT^* and Treg.*iEzh2^Y641F^* mice.

(B) Quantified frequencies of CD103^+^, TIGIT^+^, and PD1^+^ of YFP^+^RFP^+^ Treg cells in the draining LN (top row) or CNS tissues (bottom row) of Treg.*Ezh2^WT^/Ezh2^WT^* and Treg.*Ezh2^Y641F^/Ezh2^WT^* mice.

(C) Representative flow cytometry plot of the frequency of *Ezh2^WT^* and *Ezh2^Y641F^* Treg cells prior to adoptive transfer.

(D-E) Representative flow cytometry plots and quantified frequencies of *Ezh2^WT^* and *Ezh2^Y641F^* Treg cells recovered from the spleen parenchyma (D) and spleen vasculature (E) after adoptive transfer.

Data are mean ± SEM (n=5-13 mice per group pooled from two or three independent experiments); two-way repeated measured ANOVA (A) or unpaired two-tailed Student’s t-test (B,C,E,F) and were used; *p < 0.05, **p < 0.01, ***p < 0.001, and ****p<0.0001 (only statistically significant differences are noted and non-significant data is not indicated).

**Figure S6. Global redistribution of H3K27me3 in *Ezh2^Y641F^* Treg cells.**

(A) Top 10 mammalian phenotype ontology terms associated with genes containing decreased H3K27me3 modifications in the promoter regions of naïve *Ezh2^WT^* versus *Ezh2^Y641F^* Treg cells.

(B) Top 10 mammalian phenotype ontology terms associated with genes containing decreased H3K27me3 modifications in the promoter regions of activated *Ezh2^WT^* versus *Ezh2^Y641F^* Treg cells.

(C) H3K27me3 modification tracks in naïve *Ezh2^WT^* versus *Ezh2^Y641F^* Treg cells and IgG control, as well as representative chromatin accessibility tracks from ATAC-seq data, for genomic regions surrounding the *Il2ra* and *Tigit* loci and labeled as in Figure 6G.

(D) Strategy to overlap H3K27me3 segments with ATAC-seq peaks.

(E) Frequency of H3K27me3 segments overlapping with ATAC-seq peaks separated by each quartile. Number of ATAC-seq peaks overlapping with H3K27me3 segments (2924 potential H3K27me3 segments/quartile) are indicated above each bar.

Genomic data are obtained from three biological replicates. Hypergeometric test after multiple testing correction (E) was used; *p < 0.05, **p < 0.01, ***p < 0.001, and ****p<0.0001 (only statistically significant differences are noted and non-significant data is not indicated).

**Figure S7. *Ezh2^Y641F^* Treg cells have H3K37me3-associated gene expression features of Treg cell and activated T cell phenotypes.**

(A) Number of domains and domain coverage (bp) identified in naïve or activated, *Ezh2^WT^* and *Ezh2^Y641F^* Treg cells.

(B) Principal component analysis of H3K27me3 enrichment across all segments in CL1 and CL2.

(C) Bar graph illustrating the score of each sample in PC1 from the PCA depicted in Figure S7B.

(D) Fraction of domains overlapping with genic features and heatmaps displaying the log2 ratio of over-representation of each class of annotation in each dataset (left) and cluster (right) compared to the sum of all datasets or clusters. Significant differences are indicated in bold.

(E) Heatmap displaying RNA expression of naïve *Ezh2^Y641^* and activated *Ezh2^WT^* and *Ezh2^Y641F^* Treg cells compared to naïve *Ezh2^WT^* Treg cells for genes associated with segments in each cluster as defined in Figure 7A-B and present in superset of genes (11,440) as defined in Figure 7C.

(F) Boxplots demonstrating the fold change in gene expression between activated *Ezh2^Y641F^* and *Ezh2^WT^* Treg cells (left) and activated *Ezh2^Y641F^* and naïve *Ezh2^Y641F^* Treg cells (right) of superset genes associated with H3K27me3 segments in each cluster (CL1-CL4). Statistics for left boxplot is as follows: CL1****; CL2***; CL3****; CL4****, statistics for right boxplot is as follows: CL1****; CL2****; CL3****; CL4*.

(G-H) Gene set enrichment analysis of genes downregulated or upregulated in lymph node-derived Treg cells versus conventional CD4^+^ T cells (gene sets obtained from GSE7582 and GSE7460) compared to gene expression in naïve *Ezh2^Y641F^* versus naïve *Ezh2^WT^* Treg cells (based on superset of genes)

(I) Gene set enrichment analysis of genes upregulated (left) in activated *Ezh2*-deficient versus WT Treg cells or upregulated (right) in naïve *Ezh2*-deficient versus WT Treg cells (obtained from GSE58998) compared to gene expression in naïve *Ezh2^Y641F^* versus naïve *Ezh2^WT^* Treg cells (based on superset of genes).

1. (J) Gene set enrichment analysis of genes downregulated (left) or upregulated (right) in anti-CD3/anti-CD28 co-stimulated versus anti-CD3 stimulated CD4^+^ T cells (obtained from GSE39595) compared to gene expression from naïve *Ezh2^Y641F^* versus naïve *Ezh2^WT^* Treg cells (based on superset of genes).

Genomic data are obtained from three biological replicates. Hypergeometric test after multiple testing correction (B) and Wilcoxon-signed rank tests (F) were used; *p < 0.05, **p < 0.01, ***p < 0.001, and ****p<0.0001 (only statistically significant differences are noted and non-significant data is not indicated).

