## Supplemental information for "Increased EZH2 function in regulatory T cells promotes their capacity to suppress autoimmunity by driving effector differentiation prior to activation"

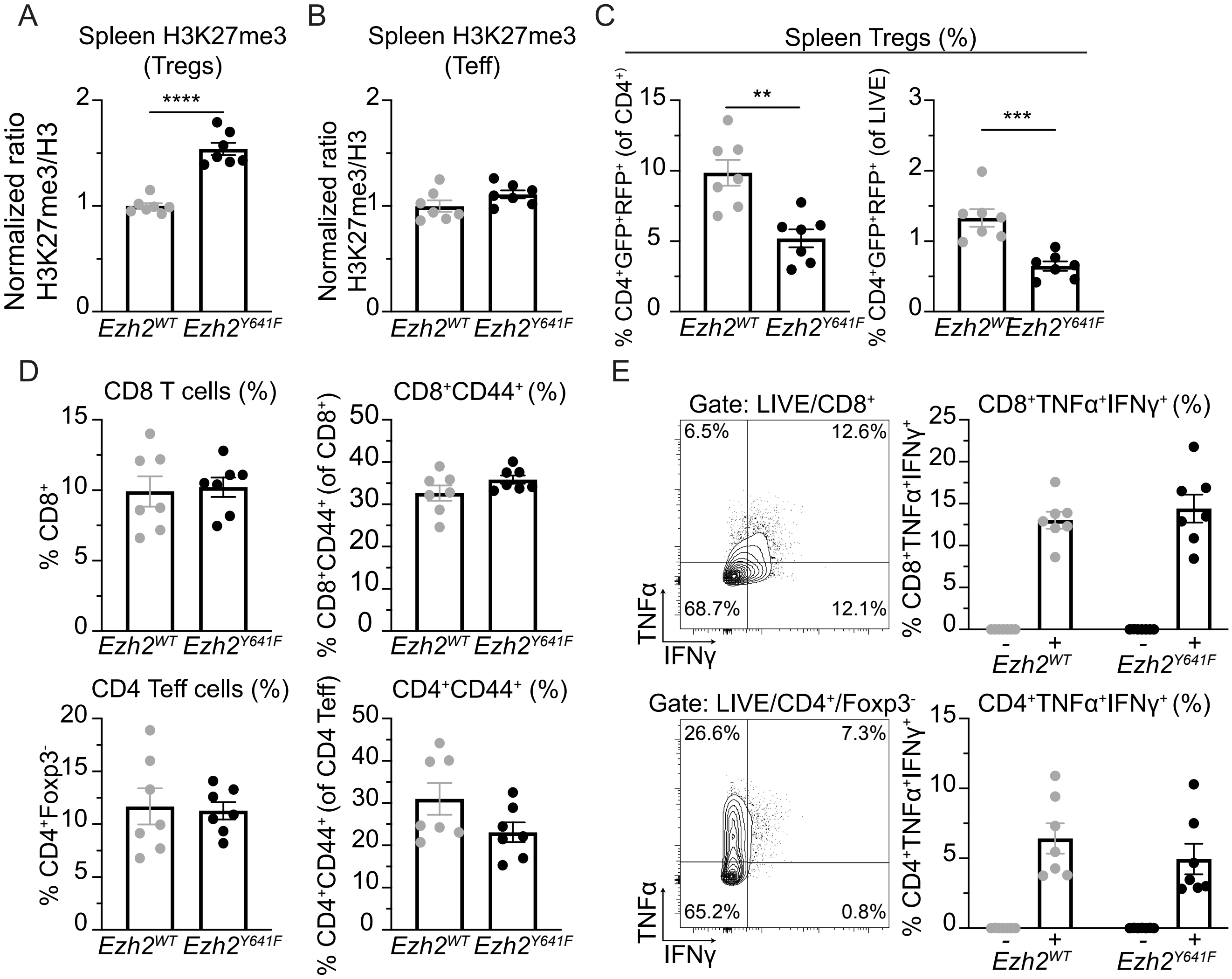


Figure S1. Peeters et al., 2024

**Figure S1. Expression of *$Ezh2^{Y641F}$* in Treg cells increases H3K27me3 levels in the spleen.**

(A-B) Normalized H3K27me3/Histone H3 levels of $CD4^{+}Foxp3^{+}$ (A) or $CD4^{+}Foxp3^{-}$ (Teff) (B) cells in spleen of Treg.$Ezh2^{WT}$ and Treg.$Ezh2^{Y641F}$ mice.

(C) Frequency of $CD4^{+}GFP^{+}RFP^{+}$ cells of $CD4^{+}$ and LIVE cells in spleen of Treg.$Ezh2^{WT}$ and Treg.$Ezh2^{Y641F}$ mice.

(D) Frequency of $CD8^{+}$, $CD8^{+}CD44^{+}$, $CD4^{+}$ Teff, and $CD4^{+}CD44^{+}$ Teff in spleen of Treg.$Ezh2^{WT}$ and Treg.$Ezh2^{Y641F}$ mice.

(E) Representative flow cytometry plots and quantified frequencies of $TNF\alpha^{+}IFN\gamma^{+}$ of $CD8^{+}$ and $CD4^{+}$ Teff cells in the spleen of Treg.$Ezh2^{WT}$ and Treg.$Ezh2^{Y641F}$ mice with (+) or without (-) *in vitro* stimulation with PMA and ionomycin.

Data are mean ± SEM and representative of at least two independent experiments (n=12-13 mice total per genotype). Unpaired two-tailed Student's t-test; $p < 0.05$, $^{**}p < 0.01$, $^{***}p < 0.001$, and $^{****}p<0.0001$ (only statistically significant differences are noted and non-significant data is not indicated).

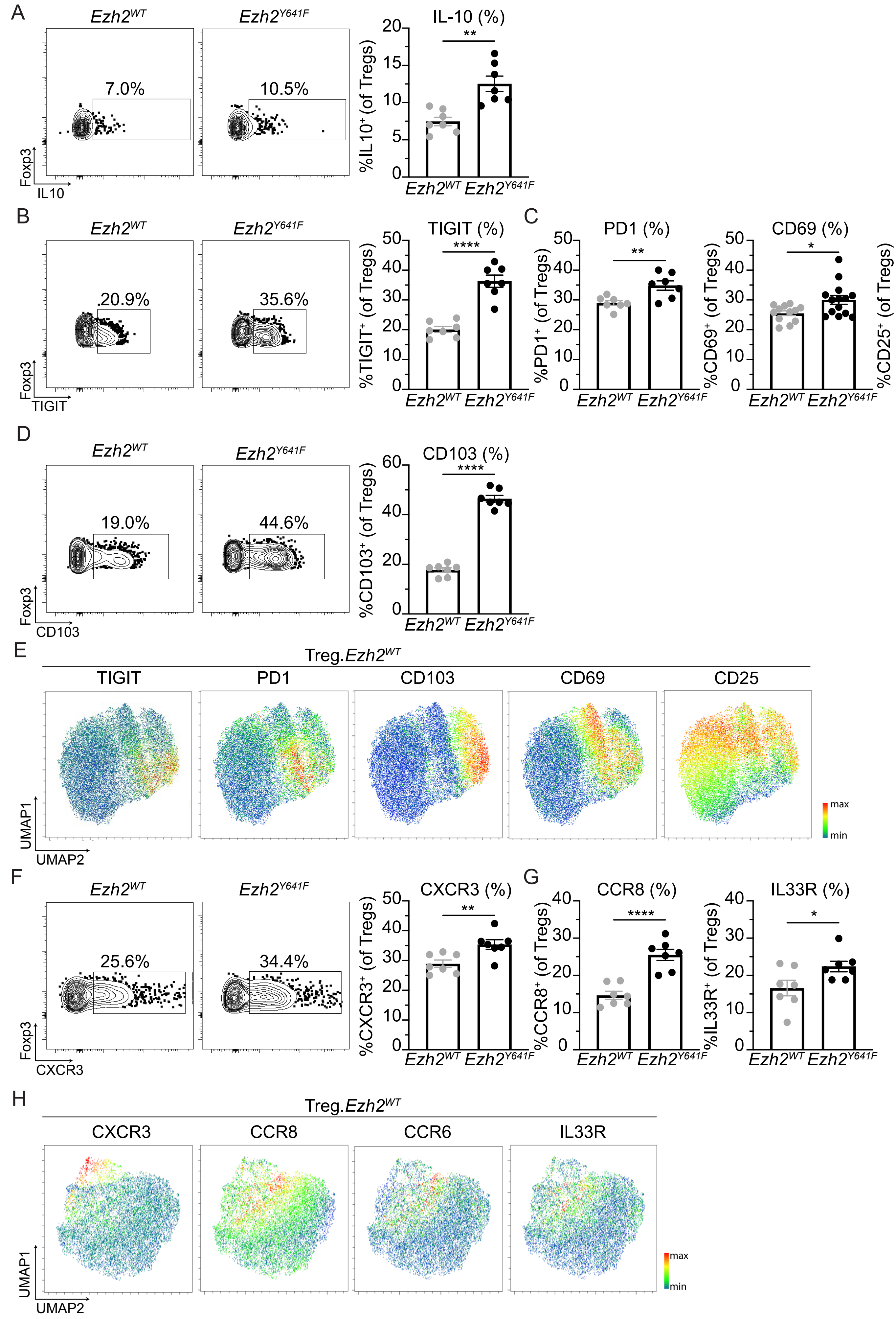


Figure S2. Peeters et al., 2024

**Figure S2.** ***Ezh2****$^{Y641F}$* **Treg cells exhibit increased activation and effector differentiation in the spleen.**

(A) Representative flow cytometry plots and quantified frequencies of IL-10$^{+}$ Treg cells from spleen of Treg.*Ezh2*$^{WT}$ and Treg.*Ezh2*$^{Y641F}$ mice.

(B) Representative flow cytometry plots and quantified frequencies of TIGIT$^{+}$ Treg cells in spleen of Treg.*Ezh2*$^{WT}$ and Treg.*Ezh2*$^{Y641F}$ mice.

(C) Quantified frequencies of PD1$^{+}$, CD69$^{+}$ and CD25$^{+}$ Treg cells in spleen of Treg.*Ezh2*$^{WT}$ and Treg.*Ezh2*$^{Y641F}$ mice.

(D) Representative flow cytometry plots and quantified frequencies of CD103$^{+}$ of Treg cells in spleen of Treg.*Ezh2*$^{WT}$ and Treg.*Ezh2*$^{Y641F}$ mice.

(E) UMAP analysis demonstrating the expression of CD103, TIGIT, PD1, CD69, and CD25 in Treg cells from LN of Treg.*Ezh2*$^{WT}$ mice.

(F) Representative flow cytometry plots and quantified frequencies of CXCR3$^{+}$ in spleen of Treg.*Ezh2*$^{WT}$ and Treg.*Ezh2*$^{Y641F}$ mice.

(G) Quantified frequencies of CCR8$^{+}$ and IL33R$^{+}$ Treg cells in spleen of Treg.*Ezh2*$^{WT}$ and Treg.*Ezh2*$^{Y641F}$ mice.

(H) UMAP analysis demonstrating the expression of CXCR3, CCR8, CCR6, and IL33R in Treg cells from LN of Treg.*Ezh2*$^{WF}$ mice.

Data are mean ± SEM and representative of at least two independent experiments (n=12-13 mice total per genotype). Unpaired two-tailed Student's t-test; *p < 0.05, **p < 0.01, ***p < 0.001, and ****p<0.0001 (only statistically significant differences are noted and non-significant data is not indicated).

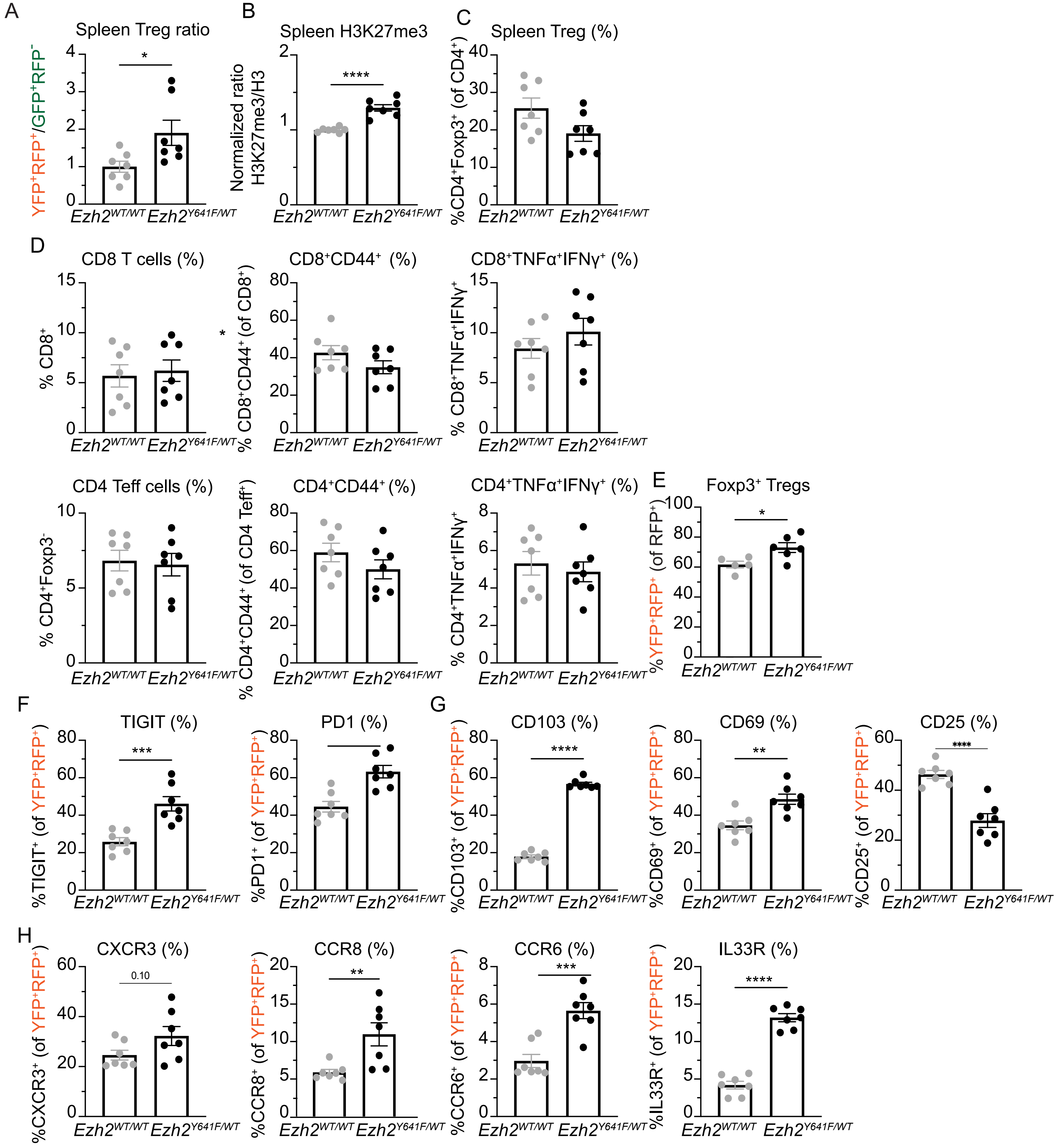


Figure S3. Peeters et al., 2024

**Figure S3. *Ezh2$^{Y641F}$* Treg cells have a competitive advantage over WT Treg cells in the spleen.**

(A) Normalized ratio $YFP^{+}RFP^{+}/GFP^{+}RFP^{-}$ Treg cells in the spleen of Treg.*Ezh2$^{WT}$/Ezh2$^{WT}$* and Treg.*Ezh2$^{Y641F}$/Ezh2$^{WT}$* female mosaic mice.

(B) Normalized H3K27me3/Histone H3 levels of $CD4^{+}Foxp3^{+}$ Treg cells in the spleen of Treg.*Ezh2$^{WT}$/Ezh2$^{WT}$* compared to Treg.*Ezh2$^{Y641}$/Ezh2$^{WT}$* mice.

(C) Frequency of $CD4^{+}Foxp3^{+}$ cells in spleen of Treg.*Ezh2$^{WT}$/Ezh2$^{WT}$* and Treg.*Ezh2$^{Y641F}$/Ezh2$^{WT}$* mice.

(D) Quantified frequencies of total $CD8^{+}$ T cells, $CD44^{+}CD8^{+}$ T cells, and $TNF\alpha^{+}IFN\gamma^{+}CD8^{+}$ T cells (with PMA/Iono restimulation) (top row) and total $CD4^{+}$ Teff cells, $CD44^{+}CD4^{+}$ Teff cells, and $TNF\alpha^{+}IFN\gamma^{+}CD4^{+}$ Teff cells (with PMA/Iono restimulation) (bottom row) in spleen of Treg.*Ezh2$^{WT}$/Ezh2$^{WT}$* and Treg.*Ezh2$^{Y641F}$/Ezh2$^{WT}$* mice.

(E) Frequency of $YFP^{+}RFP^{+}$ Treg cells (as a percentage of $CD4^{+}RFP^{+}$ cells) in spleen of Treg.*Ezh2$^{WT}$/Ezh2$^{WT}$* and Treg.*Ezh2$^{Y641F}$/Ezh2$^{WT}$* mice.

(F) Quantified frequencies of $TIGIT^{+}$ and $PD1^{+}$ of $YFP^{+}RFP^{+}$ Treg cells in the spleen of Treg.*Ezh2$^{WT}$/Ezh2$^{WT}$* and Treg.*Ezh2$^{Y641F}$/Ezh2$^{WT}$* mice.

(G) Quantified frequencies of $CD103^{+}$, $CD69^{+}$, $CD25^{+}$ of $YFP^{+}RFP^{+}$ Treg cells in the spleen of Treg.*Ezh2$^{WT}$/Ezh2$^{WT}$* and Treg.*Ezh2$^{Y641F}$/Ezh2$^{WT}$* mice.

(H) Quantified frequencies of $CXCR3^{+}$, $CCR8^{+}$, $CCR6^{+}$, and $IL33R^{+}$ of $YFP^{+}RFP^{+}$ Treg cells in the spleen of Treg.*Ezh2$^{WT}$/Ezh2$^{WT}$* and Treg.*Ezh2$^{Y641F}$/Ezh2$^{WT}$* mice.

Data are mean ± SEM and representative of at least two independent experiments (n=8-11 mice total per genotype). Unpaired two-tailed Student's t-test; $*p < 0.05$, $**p < 0.01$, $***p < 0.001$, and $****p<0.0001$ (only statistically significant differences are noted and non-significant data is not indicated).

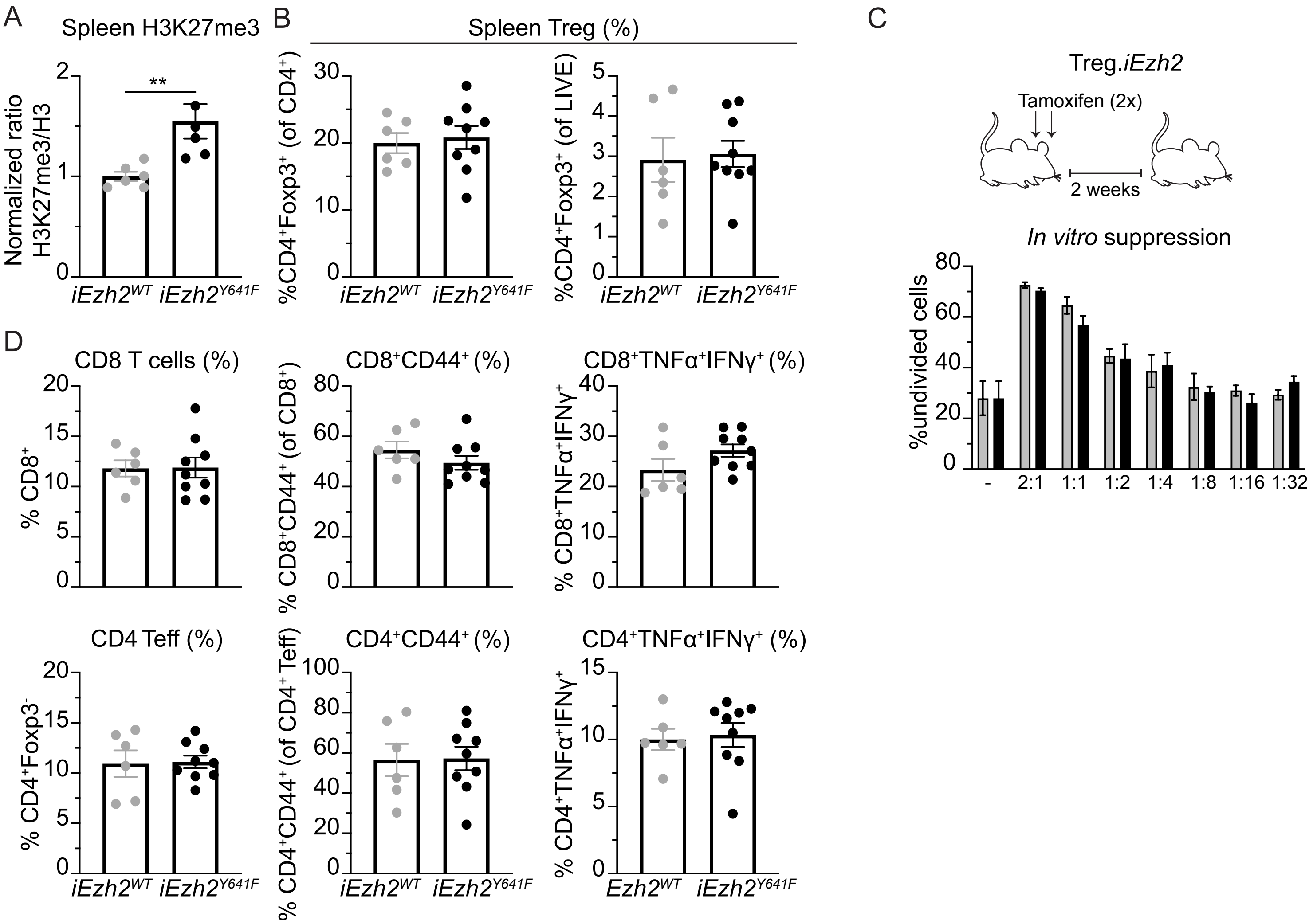


Figure S4. Peeters et al., 2024

**Figure S4. Acute induction of *Ezh2$^{Y641F}$* does not immediately impact T effector cell homeostasis.**

(A) Normalized H3K27me3/Histone H3 levels of $CD4^+Foxp3^+$ Treg cells in the spleen of *Ezh2$^{Y641F}$*-inducible Treg.*iEzh2$^{WT}$* compared to Treg.*iEzh2$^{Y641F}$* mice two weeks after tamoxifen treatment.

(B) Quantified frequencies of $CD4^+Foxp3^+$ Treg cells in the spleen of *Ezh2$^{Y641F}$*-inducible Treg.*iEzh2$^{WT}$* and Treg.*iEzh2$^{Y641F}$* mice two weeks after tamoxifen treatment.

(C) Frequencies of undivided $CD4^+$ T effector cells after co-culture with indicated ratio of Treg/Teff cells (- indicates no Treg cells) (one of three biological replicates shown). Treg cells were FACS purified from *Ezh2$^{Y641F}$*-inducible Treg.*iEzh2$^{WT}$* and Treg.*iEzh2$^{Y641F}$* mice two weeks after tamoxifen treatment.

(D) Quantified frequencies of total $CD8^+$ T cells, $CD44^+CD8^+$ T cells, and $TNF\alpha^+IFN\gamma^+CD8^+$ T cells (with PMA/Iono restimulation) (top row) and total $CD4^+$ Teff cells, $CD44^+CD4^+$ Teff cells, and $TNF\alpha^+IFN\gamma^+CD4^+$ Teff cells (with PMA/Iono restimulation) (bottom row) in spleen of Treg.*iEzh2$^{WT}$* and Treg.*iEzh2$^{Y641F}$* mice two weeks after tamoxifen treatment.

Data are mean ± SEM and representative of at least two independent experiments (n=10-14 mice total per genotype), unless noted otherwise. Unpaired two-tailed Student's t-test; *$p < 0.05$, **$p < 0.01$, ***$p < 0.001$, and ****$p<0.0001$ (only statistically significant differences are noted and non-significant data is not indicated).

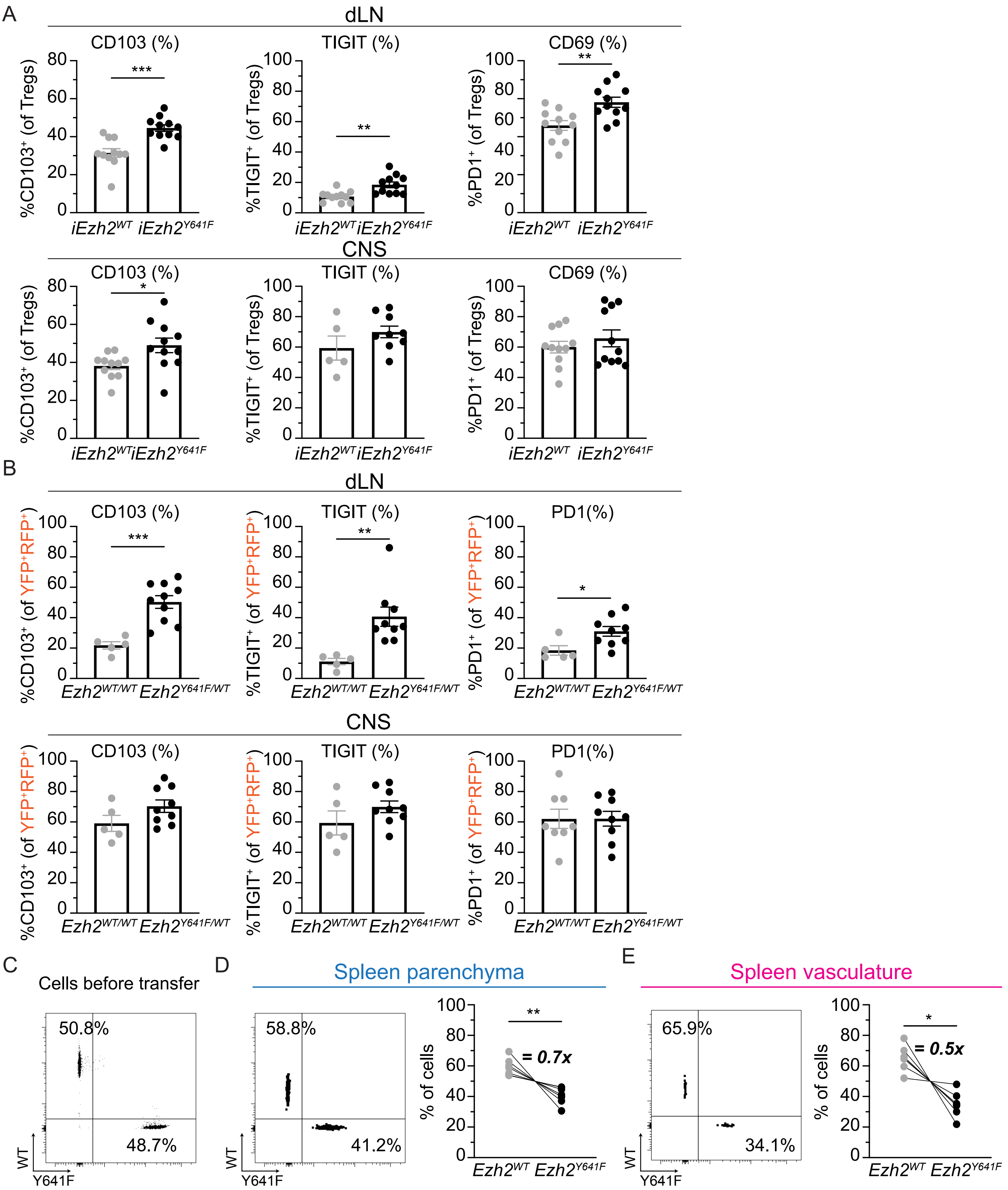


Figure S5. Peeters et al., 2024

**Figure S5. Enhanced effector differentiation of *Ezh2*$^{Y641F}$ in LN is associated with increased migration to organ tissues.**

(A) Quantified frequencies of CD103$^{+}$, TIGIT$^{+}$, and CD69$^{+}$ Treg cells within the draining LN (top row) or CNS tissues (bottom row) of Treg.*iEzh2*$^{WT}$ and Treg.*iEzh2*$^{Y641F}$ mice.

(B) Quantified frequencies of CD103$^{+}$, TIGIT$^{+}$, and PD1$^{+}$ of YFP$^{+}$RFP$^{+}$ Treg cells in the draining LN (top row) or CNS tissues (bottom row) of Treg.*Ezh2*$^{WT}$/*Ezh2*$^{WT}$ and Treg.*Ezh2*$^{Y641F}$/*Ezh2*$^{WT}$ mice.

(C) Representative flow cytometry plot of the frequency of *Ezh2*$^{WT}$ and *Ezh2*$^{Y641F}$ Treg cells prior to adoptive transfer.

(D-E) Representative flow cytometry plots and quantified frequencies of *Ezh2*$^{WT}$ and *Ezh2*$^{Y641F}$ Treg cells recovered from the spleen parenchyma (D) and spleen vasculature (E) after adoptive transfer.

Data are mean ± SEM (n=5-13 mice per group pooled from two or three independent experiments); two-way repeated measured ANOVA (A) or unpaired two-tailed Student's t-test (B,C,E,F) and were used; *$p < 0.05$, **$p < 0.01$, ***$p < 0.001$, and ****$p<0.0001$ (only statistically significant differences are noted and non-significant data is not indicated).

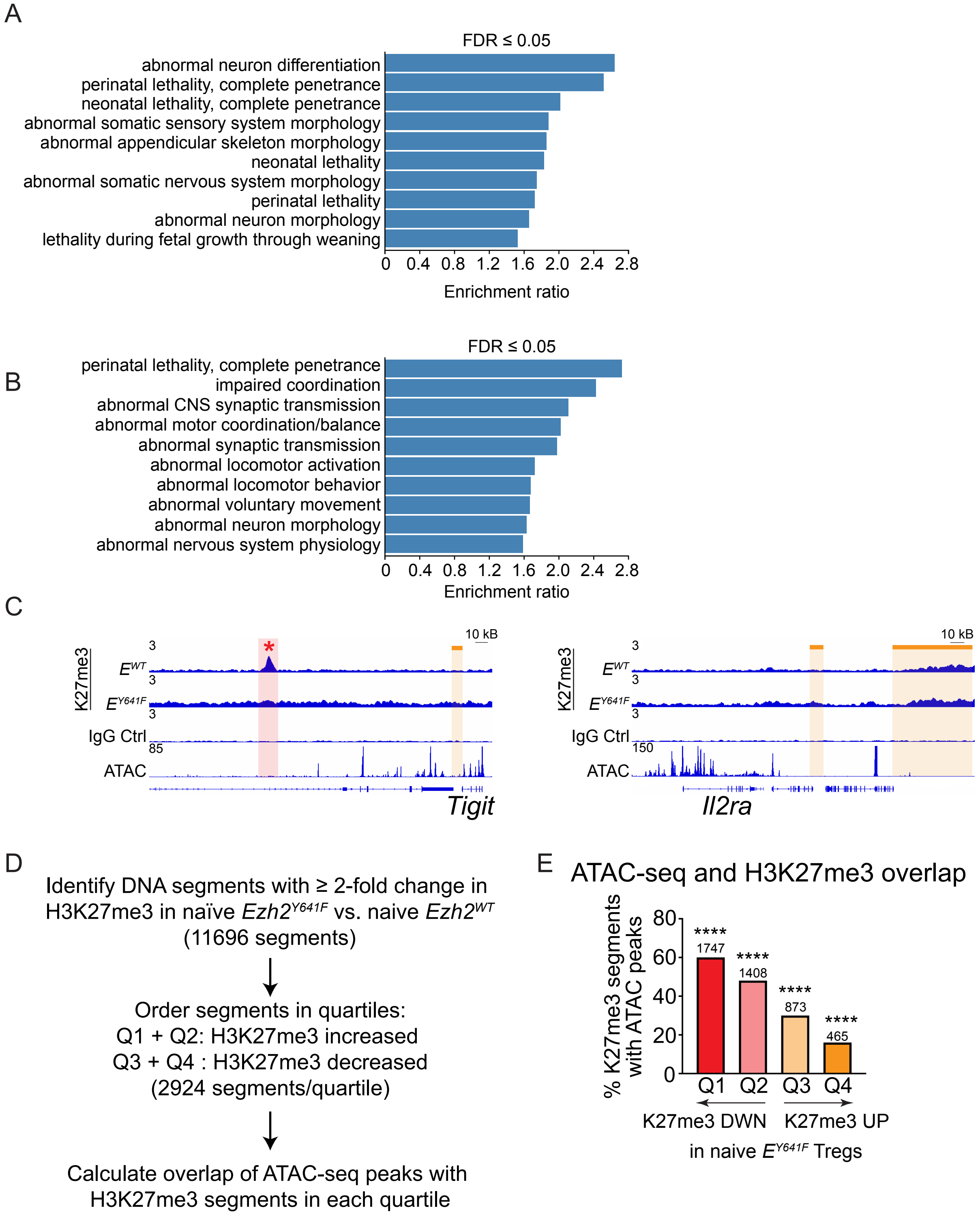


Figure S6. Peeters et al., 2024

**Figure S6. Global redistribution of H3K27me3 in *Ezh2*$^{Y641F}$ Treg cells.**

(A) Top 10 mammalian phenotype ontology terms associated with genes containing decreased H3K27me3 modifications in the promoter regions of naïve *Ezh2*$^{WT}$ versus *Ezh2*$^{Y641F}$ Treg cells.

(B) Top 10 mammalian phenotype ontology terms associated with genes containing decreased H3K27me3 modifications in the promoter regions of activated *Ezh2*$^{WT}$ versus *Ezh2*$^{Y641F}$ Treg cells.

(C) H3K27me3 modification tracks in naïve *Ezh2*$^{WT}$ versus *Ezh2*$^{Y641F}$ Treg cells and IgG control, as well as representative chromatin accessibility tracks from ATAC-seq data, for genomic regions surrounding the *Il2ra* and *Tigit* loci and labeled as in Figure 6G.

(D) Strategy to overlap H3K27me3 segments with ATAC-seq peaks.

(E) Frequency of H3K27me3 segments overlapping with ATAC-seq peaks separated by each quartile. Number of ATAC-seq peaks overlapping with H3K27me3 segments (2924 potential H3K27me3 segments/quartile) are indicated above each bar.

Genomic data are obtained from three biological replicates. Hypergeometric test after multiple testing correction (E) was used; $*p < 0.05$, $**p < 0.01$, $***p < 0.001$, and $****p<0.0001$ (only statistically significant differences are noted and non-significant data is not indicated).

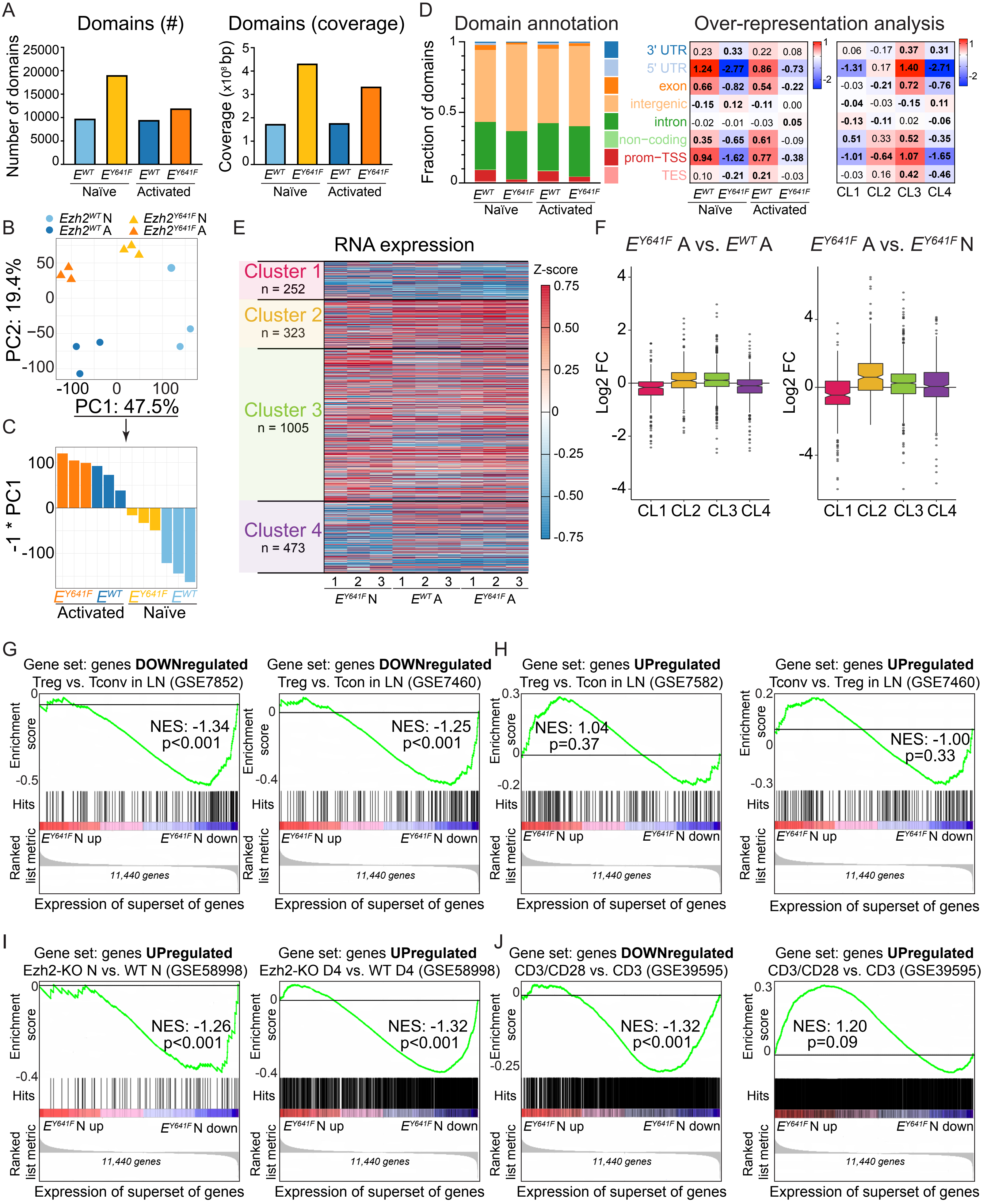


Figure S7. Peeters et al., 2024

**Figure S7. *Ezh2$^{Y641F}$* Treg cells have H3K37me3-associated gene expression features of Treg cell and activated T cell phenotypes.**

(A) Number of domains and domain coverage (bp) identified in naïve or activated, *Ezh2$^{WT}$* and *Ezh2$^{Y641F}$* Treg cells.

(B) Principal component analysis of H3K27me3 enrichment across all segments in CL1 and CL2.

(C) Bar graph illustrating the score of each sample in PC1 from the PCA depicted in Figure S7B.

(D) Fraction of domains overlapping with genic features and heatmaps displaying the log2 ratio of over-representation of each class of annotation in each dataset (left) and cluster (right) compared to the sum of all datasets or clusters. Significant differences are indicated in bold.

(E) Heatmap displaying RNA expression of naïve *Ezh2$^{Y641}$* and activated *Ezh2$^{WT}$* and *Ezh2$^{Y641F}$* Treg cells compared to naïve *Ezh2$^{WT}$* Treg cells for genes associated with segments in each cluster as defined in Figure 7A-B and present in superset of genes (11,440) as defined in Figure 7C.

(F) Boxplots demonstrating the fold change in gene expression between activated *Ezh2$^{Y641F}$* and *Ezh2$^{WT}$* Treg cells (left) and activated *Ezh2$^{Y641F}$* and naïve *Ezh2$^{Y641F}$* Treg cells (right) of superset genes associated with H3K27me3 segments in each cluster (CL1-CL4). Statistics for left boxplot is as follows: CL1****; CL2***; CL3****; CL4****, statistics for right boxplot is as follows: CL1****; CL2****; CL3****; CL4*.

(G-H) Gene set enrichment analysis of genes downregulated or upregulated in lymph node-derived Treg cells versus conventional CD4$^{+}$ T cells (gene sets obtained from GSE7582 and GSE7460) compared to gene expression in naïve *Ezh2$^{Y641F}$* versus naïve *Ezh2$^{WT}$* Treg cells (based on superset of genes)

(I) Gene set enrichment analysis of genes upregulated (left) in activated *Ezh2*-deficient versus WT Treg cells or upregulated (right) in naïve *Ezh2*-deficient versus WT Treg cells (obtained from GSE58998) compared to gene expression in naïve *Ezh2$^{Y641F}$* versus naïve *Ezh2$^{WT}$* Treg cells (based on superset of genes).

(J) Gene set enrichment analysis of genes downregulated (left) or upregulated (right) in anti-CD3/anti-CD28 co-stimulated versus anti-CD3 stimulated CD4$^{+}$ T cells (obtained from GSE39595) compared to gene expression from naïve *Ezh2$^{Y641F}$* versus naïve *Ezh2$^{WT}$* Treg cells (based on superset of genes).

Genomic data are obtained from three biological replicates. Hypergeometric test after multiple testing correction (B) and Wilcoxon-signed rank tests (F) were used; *p < 0.05, **p < 0.01, ***p < 0.001, and ****p<0.0001 (only statistically significant differences are noted and non-significant data is not indicated).
